# The influence of different sugars and sugar alcohol, and cultivars in the osmotically dehydrated peach fruit. Analysis of mass transfer parameters and the variation in the organoleptic properties, bioactive and nutritive compounds, and pro-health properties in dried peaches

**DOI:** 10.1101/2025.02.07.637111

**Authors:** Lara Salvañal, Julieta Gabilondo, Graciela Corbino, Claudio O. Budde, María Valeria Lara

## Abstract

Peach is a great-taste fruit, source of nutrients and bioactive compounds. It is a seasonal fruit that cannot be stored for prolonged periods of time. To increase its consumption over the year, promote its intake as a snack, utilize it in the industry, process the overproduced peach and to add value to the fruit, osmotic dehydration (OD) followed by hot-air drying (OD+D) is a suitable process. Here, we compared the use of three different osmolytes (glucose, sorbitol or sucrose) and four peach cultivars (Flordaking, Goldprince, Elegant Lady and Dixiland) exposed to OD+D to study the effects on organoleptic, nutritional and nutraceutical properties of the peach. OD+D slices were compared against fresh fruit exclusively exposed to hot-air treatment (D). A moisture content of 17.5% was obtained in OD+D and D peaches. Moisture content, water loss, solute gain and colour depended on the cultivar. Higher metabolite and antioxidant capacity retention was observed for OD+D peach slices compared to D peaches, minimizing adverse changes such as carotenoid decrease typical of D and even enhancing some features such as tannin content. Moreover, OD+D improved the inhibition of protease, α-amylase and α-amyloglucosidase, key enzymes involved in inflammatory processes and hyperglycemia management. The dehydration processes modified the antimicrobial capacity of peach slices, with responses varying depending on the cultivar and obtaining better inhibitions than in D. Finally, although mineral composition was altered in OD+D due to leaking to the solution, the addition of calcium in the hyperosmotic solution resulted in calcium enriched dried peaches. Collectively, the results obtained reveal the influence of the cultivar on the properties of the dried fruit and highlight the benefits of the osmotic pretreatments prior to conventional drying.

## 1. Introduction

Peach is among the most important fruit species from temperate climate produced for direct consumption and for industrialization. Peach fruit has global production of 24.5 million tons on a cultivated area of around 1.5 million ha in 2020 (FAOSTAT 2022). It is consumed worldwide because of its color, flavor and aroma characteristics. In addition to its great-taste, it is a source of nutrients and bioactive compounds (Lara et al., 2020). The peach is highly perishable; it ripens at room temperature and due to its high moisture, peach deteriorates in a relatively short time (Crisosto et al., 1999; Crisosto et al., 2006; Lurie & Crisosto, 2005). Besides low temperature storage (0-5°C) is used to extend its shelf-life and to avoid fruit decay during commercialization it can conduct to the initiation of the chilling injury disorder (Pedreschi & Lurie, 2015). In any case, the storage time is limited, with some cultivars (cvs) reaching 5 weeks. Therefore, it is important the use of conservation techniques that offer products similar to fresh ones while retaining their nutritional properties. Additionally, it is essential to identify alternative uses for fruits that do not enter the fresh market, either because they do not meet established quality standards and are considered waste, or due to seasonal overproduction. The dehydration of fresh fruits is an alternative destination for the fruit, and dehydrated peaches constitute a healthy product that can reach the consumer or be used in the food processing industry. Fruit snacks can be obtained through dehydration and are stable foods with low water activity offering a very pleasant flavor and texture. Dehydrated fruits can be consumed at any time of the day; they do not need cooking and are available all year round. Moreover, dehydration techniques not only extend the shelf life of foods without the need for refrigeration but also lead to reduction in costs in food related to food storage, packaging, transportation and distribution and it increase the economic value of the fruit commodity. In addition, dehydrated fruits participate in a larger segment of the market: the food processing industry, where they are used as raw materials in the production of cereal bars, yogurts, ice creams and baked goods.

In recent decades, consumers have become more aware of the relationship between diet and disease. Nowadays, there is a broad consensus that increased consumption of fruits and vegetables contributes to improved health and well-being by reducing the risk of diseases such as cardiovascular diseases and some types of cancer (Hung et al., 2004; Riboli et al., 2003).

In this respect, peach fruit has an important therapeutic and nutritional value (Kurz et al., 2008). Peach is one of the most widely consumed fruits in several European countries, especially those with the Mediterranean diet (Konopacka et al., 2010; SauraCalixto & Goñi, 2006). Epidemiological studies suggest that the Mediterranean diet, which is partially based on fruit consumption, may play a role in the prevention of several diseases (Sofi et al., 2008). The consumption of peaches can reduce the production of reactive oxygen species (ROS) in human plasma and provides protection against different diseases (Tsantili et al., 2010) due to the presence of compounds with significant anti-free radical power properties (Cevallos-Casals et al., 2006). Together, the need to incorporate more fruits in the daily diet drives the agroindustry to adopt conservation techniques that deliver products resembling fresh fruit.

Osmotic dehydration (OD) is technique applied before conventional hot air drying (D) to improve the sensorial, nutritional and functional quality of the product and to reduce time and energy during the processing and to decrease the negative effects of heat on food (Ahmed et al., 2016). OD can also be applied before freezing (Ramallo & Mascheroni, 2010), frying (Taiwo & Baik, 2007) or vacuum drying (Ferrari et al., 2011). OD partially removes water from the foodstuff by immersing it in a hypertonic solution with one or more solutes (Ponting et al., 1966). As the high osmolarity of the solution drives the water out of the food, solutes from the solution enter the tissue. In addition, some solutes from the food can migrate into the osmotic solution (Raoult –Wack, 1994). The kinetics of OD depends on the solution composition and concentration, fruit: solution ratio, the parameters of the process such as temperature, time and agitation as well as food properties including size and geometry, tissue characteristics and fruit ripeness level (Chiralt & Fito, 2003). Most common osmotic agents used include sucrose, glucose, oligosaccharides, sorbitol, glycerol, glucose with corn syrup, and salts (sodium chloride).

OD has been successfully applied to fruits like apple (Amami et al., 2006), mango (Zhao et al., 2013), pineapple (Beristain et al., 1990), pear (Park et al., 2002) and tomato (Giannakourou et al., 2020), vegetables like carrot (Peng et al., 2019) and bamboo shoots (Badwaik et al., 2013), meat (Dimakopoulou-Papazoglou & Katsanidis, 2020), fish and mollusks (Lemus-Mondaca et al., 2014).

Although OD of peach has recently received attention as a pre-treatment to conventional drying, most efforts have been dedicated to optimizing the process conditions for OD, exploring the mass transport, texture properties, sugar content and sensory attributes of the dehydrated peach (Dhillon et al., 2022; Giangiecome et al., 1987; Marconi Germer et al., 2010; Sahari et al., 2006; Wang et al., 2023a,b; Yadav and Singh, 2014). Nevertheless, research exploring the effect of OD+D on nutritional properties and on other beneficial characteristics of the peach fruit such as the inhibition of microbial growth or inflammatory enzymes has not been conducted. Since consumers are requesting minimally processed foods that preserve their nutritional properties, the study of the functional characteristics of dehydrated peach is of great relevance. Thus, the objective of this work was, on one hand, to compare the quality and properties of peach fruits osmotically dehydrated prior to conventional hot air drying (D) with those fruit subjected to exclusively conventional D and those of fresh fruit (F). On the other hand, we aimed to compare the use of three different osmolytes (Glucose, Sorbitol or Sucrose) and four cultivars of peach during the OD to establish optimum solution and cultivar for OD. As a first step towards this objective, the peach cultivars Flordaking (FD), Goldprince (GP), Elegant Lady (EL) and Dixiland (DX) were characterized regarding their organoleptic, nutritional and nutraceutical properties.

## 2. Materials and methods

### 2.1. Fruit material and treatments

Assays were developed in four cultivars of *Prunus persica* (L.) Batsch: the earlier harvested, Flordaking (FD) and Goldprince (GP) and the mid-season Elegant Lady (EL) and Dixiland (DX). Fruits were harvested from November to January, depending on the cultivar, in the 2019-2020, 2020-2021 and 2021-2022 seasons in a commercial orchard (ARTIGUES - Parana Basin Fruit S.R.L) located in San Pedro, Buenos Aires, Argentina (33°40′46″S 59°40′01″O)

Peach fruit was washed with 200 ppm chlorinated water at pH 6.8, for 2 min, rinsed with tap water and sliced with skin included into 2 mm thick slices using a ceramic knife. Then, slices were submerged in 20 ppm chlorinated water for 2 min, drained, immersed in an anti-browning solution for 2 min (1% (w/v) ascorbic acid (AsA) and 0.5% (w/v) citric acid) and drained for another 15 min. These samples constituted the Fresh fruit (F). A fraction was analyzed as F and another portion was placed in an oven at 58 °C with air circulation until a moisture content smaller than 18% was reached (D) or subjected to osmotic dehydration (OD) as following described. Sucrose (suc), glucose (glc) and sorbitol (sor) hyperosmotic solutions (HS, 46-47 °Brix) containing 2% (w/v) CaCl2 were used for OD. One hundred and fifty grams of fresh peach slices were placed in screw-cap jars with 1.5 L of HS and incubated at 40 °C for 3 h under orbital agitation. After incubation, slices were drained on blotting paper for 20 min. One half of this material was analyzed (OD), while the other half was dried in oven like D slices (OD+D).

After processing samples F, D, OD and OD+D were immediately used for weighting, dry matter and moisture content, mass transfer and organoleptic parameters analysis while a fraction of the samples was frozen in N2(l) and stored at -80 °C for further use. The frozen samples were homogenized in mortar with N2(l) using mortar and pestle. Homogenized powders were used for analysis.

### 2.2. Dry matter and moisture content

Water content of all samples was determined by weighing. Representative samples were weighed to record their weight (*m*) and reduced to dryness at 58°C until no further weight change occurred (dry matter, *mmd*). The *m* and *mmd* parameters were used to express the content of the different metabolites and activities evaluated in this work in term of fresh (FW) or dry weight (DW), respectively.

The moisture content (MC) was expressed in percentage and calculated as the amount of eliminated water from the samples using the following equation:

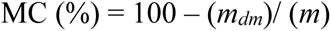

### 2.3. Mass transfer parameters

Water loss (WL), solute gain (SG) and mass loss (ML) during OD were calculated using the following equations as described in Janowicz et al. (2021):

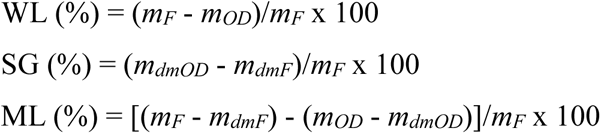

where *mF* is the initial FW of F peach slices (g), *mOD* is the FW of peach after osmotic dehydration, *mdmF* is the weight of solids (dry matter) in the F peach slices (g), and *mdmOD* is the weight of solids (dry matter) of peach slices after osmotic dehydration (g).

### 2.4. Organoleptic parameters

Colour measurements of the flesh were performed using a chromameter (Minolta CR 400, Japan) using the CIELab color system. In each slice three random measurements were conducted. L measures lightness. Negative a* values indicate greenness and positive values indicate redness. Negative b* values indicate blueness and positive values represent yellowness (Budde et al., 2006). Representative (mean of measurements) La*b* parameters were mapped to colors on a figure using the “Colorspace” package from “Rstudio”. ΔE*76 parameter was calculated to define the colour changes that are perceivable (López & Di Sarli, 2016).

Flesh firmness was evaluated in whole peaches on two paired sides of each fruit using a penetrometer (Effegi 327, Italy) with a 7.9 mm probe diameter and expressed in Kg/cm^2^.

Titratable acidity (TA) was determined in 10 g of homogenized samples in 100 g of distilled water, composed of a pool of three fruits, by titration with 0.1 M NaOH, and expressed as % malic acid.

The total soluble solids (TSS) were determined by using a refractometer (Atago N1 0–32 °C, Japan) and expressed in °Brix.

Organoleptic parameters were measured using at least fifteen fruits.

### 2.5. Sugar and sugar alcohol contents

Glucose, sucrose and sorbitol were extracted using 50 mg of pulverized peach material and 600 µl methanol for 15 min a 70 °C. Afterwards, 500 μl of cold H2Od were added. After vortexing and centrifugation at 14,000 *g* for 15 min, the supernatant was recovered and the volume was measured.

Sorbitol was measured enzymatically using sorbitol dehydrogenase from sheep (Sigma, St Louis, MO, USA, Borsani et al., 2009).

Glucose (mmol per gram of FW (GFW) or DW (GDW)) was quantified using the glucose oxidase method (Wiener lab) and the manufacturer’s instructions. For this purpose, 1 μL of a 1/100 dilution of methanolic extract was prepared in 0.5 M acetic acid-acetate buffer, pH 4.5. Absorbance was measured at 505 nm using a plate reader EPOCH2T-BioTek instruments. A calibration curve was prepared using glucose as a standard.

Sucrose was quantified by glucose measuring after hydrolysis using invertase. One microliter of 1/10 dilution of extract was incubated with 89 µl of 0.5 M acetic acid-acetate buffer, pH 4.5 and 10 µl of invertase (10 mg/ml, 30 U) from *Saccharomyces cerevisiae* (Sigma) during 40 min at 55 °C. Glucose was quantified in the samples before and after hydrolysis. The glucose produced was equivalent to sucrose hydrolysed, which was expressed as mmol per GFW or GDW.

### 2.6. Content of total phenolics

Phenolic compounds were extracted and measured using the Folin-Ciocalteau reagent (Merck) (Cantín et al., 2009). Briefly, 40 mg of pulverized material were extracted in 1 ml of acidic methanol (0.5 M HCl, 80% (v/v) CH3OH and incubated overnight (ON) at 4 °C. Supernatants were transferred to Eppendorf tubes and the obtained volume was exactly measured. One hundred and ten microliters of H2Od was added to 40 µl of extract, standard or blank (acidic methanol), followed by 150 µl of Folin–Ciocalteau reagent, mixed well and incubated at 20 °C for 3 min. Afterwards, 300 µl of sodium carbonate was added. The samples were mixed immediately and the tubes were incubated for 1 h at 20 °C in the dark. After centrifugation at maximum speed, the absorbance of the supernatants was then measured at 725 nm in an ultraviolet–visible (UV–Vis) spectrophotometer (Jenway™ 6715, Fisher Scientific, UK). Standard calibration curves, using chlorogenic acid in 80 % (v/v) CH3OH as a solvent, were prepared. The content of total phenolics (TPC) was expressed as µg chlorogenic acid equivalents (CA)/ GFW or GDW.

### 2.7. Flavonoids

Flavonoids were quantified using 50 mg of frozen pulverized peach material and 1 ml of acidic methanol (0.5 M HCl in 80% (v/v) CH3OH). After incubation ON in the dark and centrifugation, the supernatant was exactly measured. To 450 µl of the supernatant, 900 µl of chloroform were added. After centrifugation, the upper phase was recovered and the absorbance at 310 nm was recorded using a spectrometer Jenway™ 6715 (Fisher Scientific, UK) spectrophotometer. A calibration curve was done using rutin standard. Total flavonoid content was expressed as µg rutin equivalents (RE) per GFW or GDW.

### 2.8. Condensed tannins content

Tannins were quantified according to Sun et al. (1998). Tannins were extracted incubating a 70°C for 15 min 50 mg of frozen powder in the presence of 500 μl of MeOH. Condensed tannins from 25 μl of methanolic extract were mixed with 125 μl of freshly prepared solution of 1:1 of 8% (p/v) HCL (methanolic solution) and vanillin (1 % (p/v) in methanol). Depolymerized and transformed in red anthocyanidols were quantified at 500 nm using a microplate reader. Tannin content was expressed as mg of catechin equivalents per GFW or GDW.

### 2.9. Carotenoids

Were spectrophotometrically quantified according to the method of Sass-Kiss et al. (2005). To 50 mg of frozen powder that was previously lyophilized during 24 h in Lyophilizer (Liotop, Model L101, Brazil), 2 ml of a mixture of hexane, acetone and methanol (1:0.5:0.5) were added. The separation of the upper phase was facilitated by agitation for 30 min. The lower phase was mixed with 1 ml of hexane and agitated for 30 min. Both upper phases were mixed and the absorbance at 450 nm was measured in a spectrometer (Power Wave XS, BIO-TEK Instruments). Carotenoid content was calculated using β-carotene specific extinction coefficient in hexane (2592 ml. g^−1^cm^−1^) and expressed as µg per GFW or GDW.

### 2.10. Ascorbic acid quantitation

Ascorbic acid (AsA) was extracted from 50 mg of tissue using 200 μl of 5 % (p/v) H3PO4. After centrifugation for 20 min at maximum speed at 4 °C, the supernatant was exactly measured and immediately used for vitamin C analysis. AsA was measured according to Okamura (1980). Fifty microliters of extracts were incubated with 150 μl of 0.3 M acid buffer (Na2HPO4 and KH2PO4) pH 7.4 containing 5 mM EDTA, 250 μl of 10 % (p/v) Trichloroacetic Acid (TCA), 200 μl of 42 % (p/v) H3PO4, 200 μl of 4 % (p/v) α, α’-dipyridyl and 100 μl of 3% (p/v) FeCl3. The mixture was incubated at 37°C for 1 h and absorbance at 525 nm was recorded using a UV/visible spectrometer (Jenway™ 6305, Fisher Scientific, UK). H3PO4 was used as blank. A calibration curve was prepared using AsA as standard. Results were expressed as µg of AsA per GFW or GDW.

### 2.11. Antioxidant capacity (AC)

Antioxidant capacity (AC) was evaluated using the 1,1-diphenyl-2-picrylhydrazyl (DPPH) and the 2,2-azinobis(3-ethylbenzothiazoline-6-sulfonic acid) (ABTS*^+^) assays. Extracts were prepared using 50 mg of pulverized peach material and 1000 µl of 80 % (v/v) methanol and incubated overnight at 20 °C. After vortexing and centrifugation at 14,000 *g* for 15 min at 4 °C, the supernatant was recovered, and the volume was measured.

Radical scavenging activity of DPPH was assessed based on Brand-Williams et al. (1995) with modifications. First, 25 μl of extracts or blank were added to 475 μl DPPH methanolic solution (25 μg/ml). The mixture was then incubated at 20°C for 30 min in the dark. The absorbance was measured at 517 nm using a UV/visible Jenway™ 6305 (Fisher Scientific, UK) spectrophotometer.

The ABTS*^+^ assay was performed as described by Drogoudi et al. (2016). Radical ABTS*+ solution was prepared by reacting equal volumes of 14 mM ABTS and 4.9 mM potassium persulfate for 16 h at 20 °C in the dark. Then, the ABTS*^+^ solution was diluted with H2Od to an absorbance of 0.700 ± 0.005 at 734 nm. Twenty-five microliters of the methanolic extract were added to 475 μl ABTS*^+^ solution, after mixing and incubating at 30 °C for 30 min the absorbance at 734 nm was recorded using the microplate reader described above. The samples were diluted so as to give 20–80% reduction of the blank absorbance.

For both assays, a calibration curve was prepared with 6-hydroxy-2, 5, 7, 8-tetramethyl chromene-2-carboxylic acid (Trolox). AC was expressed as mmol of Trolox equivalents (TE) per GFW or GDW. Quintuplicate measurements were performed for each sample.

### 2.12. Mineral quantification

Fifty milligrams of peach fruit were completely mineralized by digestion with 3 ml 65 % (w/v) HNO3 for 5 h in a hot water bath (100°C). Ultrapure water (Millipore, France) was used to dilute 1/6 the mineral solutions. After filtering with 0.22-0.45 μm syringe filter (Sartorius) into a 15 mL falcon tube, samples were loaded into an inductively coupled plasma-mass spectrometry (ICP-MS, NexION 350X, Perkin Elmer) with a Direct Infusion (ID) sample introduction system and a quadrupole mass analyzer. The following multi-elemental calibration solutions (0.01^−1^ mg L^−1^) were used; SS-Low Level Elements ICV Stock and ILM 05.2 ICS Stock 1 (VHG Labs, Manchester, NH 03103 USA), and Multi-Element Plasma Standard Solution 4, Specpure® (Alfa AesarGmbH y Co KG, Germany). Five biological replicates were analyzed. Total mineral concentrations were expressed as µg per GFW or GDW.

### 2.13. Protein extraction and quantitation

Samples were ground in N2(l) and total protein was extracted in 100 mM Tris-HCl, pH 8.0; 1 mM EDTA; 10 mM MgCl2; 10 mM β-mercaptoethanol and 20% (v/v) glycerol. After centrifugation at 10,000 *g* for 15 min at 4°C, the supernatant was used for protein quantitation and diluted in 0.25 M Tris-HCl, pH 7.5; 2% (w/v) SDS; 0.5% (v/v) β-mercaptoethanol and 0.1% (v/v) bromophenol blue and boiled for 2 min for SDS-PAGE or precipitated with trichloroacetic acid (TCA) for proteomic analysis. Protein concentration was determined as described in Borsani et al. (2009).

### 2.14. Inhibition assays

For extract preparation, 5 g of peach were lyophilized (Lyophilizer Liotop, Model L101, Brazil), for 24 h and then pulverizing using N2(l), mortar and pestle and extracted with 50 ml of CH3OH. After mixing by agitation for 24 h at 40 °C, the mixture was centrifuged at 3,500 *g* for 20 min. The pellet was dried and resuspended in dimethyl sulfoxide (DMSO). Extracts were stored at 4 °C until further use.

#### 2.14.1. *In vitro* antimicrobial screening

Inhibition assays were based on modified Kirby–Bauer disk-diffusion method following the recommendations from the National Committee for Clinical Laboratory Standards (NCCLS, 2004). The bacteria tested were *Bacillus cereus* (ATCC 14579)*, Staphylococcus aureus* (RN4220), *Acinetobacter baumannii* (ATCC 17978), *Enterobacter faecalis* (ATCC 29212), *Salmonella typhimurium* (ATCC 14028s) and *Escherichia coli* (ATCC 25923), while the yeast tested included *Candida albicans* (ATCC 25922)*, C. tropicalis* (CCC 1311997)*, C. glabrata* (CCC 164-18)*, C. krusei* (CCC 165-18), and *C. parapsilosis* (CCC 106-18) which were available in the Mycology Reference Center (CEREMIC), National University of Rosario, Rosario, Argentina. The media used for analysis were Mueller-Hinton agar (Britania, Argentina) for bacteria and Mueller-Hinton Agar containing 2 % (w/v) Glucose and 0.5 µg/mL Methylene Blue Dye (GMB) for yeasts. Ampicillin (5 μg) and Fluconazole (10 μg) served as positive controls for antimicrobial activity while DMSO was used as negative control. Standard filter disks with a diameter of 6.0 mm were impregnated with 700 µg of peach extracts dissolved in DMSO. Plates were incubated 24 h at 37 °C for bacterial strains and for 48 h at 30 °C for the yeast strains. Diameters of the inhibition zones of microbial growth were measured. The test was performed at least in triplicate.

#### 2.14.2. Enzymes inhibition

α-amylase (AMY) inhibition assay was conducted as in Belhadj et al. (2016). Two hundred microliters of extract diluted in DMSO were pre-incubated with 500 μl of 0.02 M phosphate buffer pH 6.9 containing 0.5 mg/ml of porcine pancreatic α-amylase (700 U/mg, Sigma). To the reaction mixture, 500 μl of 1% (w/v) starch solution prepared in 0.02 M phosphate buffer pH 6.9 were added. After incubation for 10 min at 25 °C 1 ml of 1 % (w/v) dinitrosalicylic acid (DNS) was added and incubated at 100 °C for 5 min. After addition of 1 ml of H2Od, the absorbance was measured at 540 nm UV/visible Jenway™ 6715 (Fisher Scientific, UK) spectrophotometer. The inhibition of AMY activity was expressed as a percentage of the activity in the absence of the extract (100 %). Three tests were performed for each sample.

The inhibition of the α-amyloglucosidase (AMG) activity was tested by using the protocol from the enzyme manufacturer with modifications. Increasing amounts of the extract dissolved in DMSO (0 -100 mg or until the plateau was reached) were reached to 200 µl with 50 mM Potassium phosphate buffer, pH 4.6-4.8 and used to pre-incubate 2 U of α-amyloglucosidase from *Aspergillus niger* (Sigma-Aldrich) for 30 min at 55 °C. Protease reaction was started by the addition of 50 µl of 1 % (w/v) starch solution. After 10 min of incubation at 55 °C the reaction was stopped by boiling for 2 min. Glucose generated was measured using 15 µL of the reaction volume and protocol previously described.

For the protease inhibition test, protease activity was assayed according to the manufacturer protocol. Ten microliters of protease from bovine pancreas (type I, Sigma-Aldrich, 0.15 U) were pre-incubated for 30 min at 37 °C with different amounts (0-100 mg) of ethanolic extracts diluted in DMSO in a final volume of 116 µl 50 mM Potassium Phosphate buffer, pH 7.5. After the addition of 384 µl of 0.65 % (w/v) casein the reaction was incubated for 5 min at 37 °C and stopped by adding 500 µl 100 % (w/v) TCA. After an incubation on ice for 1 h min and centrifugation at 14,000 *g* for 15 min the supernatant was recovered. To 100 µl of supernatant, 50 µl of 50 mM Potassium Phosphate buffer, pH 7.5, 150 µl of Folin-Ciocalteau reagent (Merck) and 300 µL of 1 M Na2CO3 were added. After reaction at 20 °C for 1 h and centrifugation at 14,000 *g*, the absorbance of the supernatant was measured at 660 nm using a UV/visible Jenway™ 6305 (Fisher Scientific, UK) spectrophotometer. The reaction in the absence of extract was used as control. The inhibition was calculated as follows:

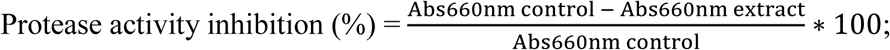

where Abs 660nm control and Abs 660 extract, correspond to the measurements of the reactions in the absence and in the presence of the peach extract, respectively.

For each enzyme, the percentage of inhibition was graph *vs* the amount of the extract. Inhibition curves were fitted to sigmoidal or hyperbolic equations using SigmaPlot and used to estimate the IC50 and maximum inhibition (Imax) of each extract.

### 2.15. Data and statistical analysis

Statistical analyses were performed using SigmaStat12.0 (SYSTAT Software) and “Agricolae” Package from Rstudio. Values are expressed as means ± SD of at least five independent replicates. Normality and homogeneity of variances were calculated. When possible, one-way analysis of variance (ANOVA) was performed, followed by Tukey’s test (α=0.05) for pairwise multiple comparisons when differences among samples were found. Kruskal-Wallis test was used when data did not pass normality tests. Two-way ANOVA was performed using the following model: cultivar (CV) + sugar (S) + interaction (CV*S) + error. When the equal variance test failed, permutational multivariate analysis of variance (PERMANOVA) was applied. Variation in the experiment was considered significant when the *p*-value was less than 0.05.

A heat map was created and hierarchical clustering analysis (HCA) using Average linkage clustering and Pearson correlation analysis (*p*-value <0.05) was conducted with MultiExperiment Viewer (MeV v4.9.0, https://webmev.tm4.org/, Saeed et al., 2003) software.

## 3. Results and discussion

### 3.1. Cultivars characterization

Four cultivars (Flordaking (FD), Goldprince (GP), Elegant Lady (EL) and Dixiland (DX)) of yellow flesh peach fruit exhibiting different postharvest dates were characterized and used in this work. Fruits were harvested at a firmness ranging between 3.91 and 5.52 Kg/cm (Fig. 1A). Organoleptic profiling revealed that pulp coloration slightly varied among cultivars, with EL exhibiting the highest reddish component (a*) and DX the highest yellowish component (b*). Regarding titratable acidity (TA), FD exhibited the highest percentage (%) of malic acid content (0.92 ± 0.05). Total soluble solids (TSS) ranged from 10.59 to 15.95 °Brix (Fig. 1A), with the earlier cvs showing the lowest values and different from the mid-season fruits.

**Fig. 1.**
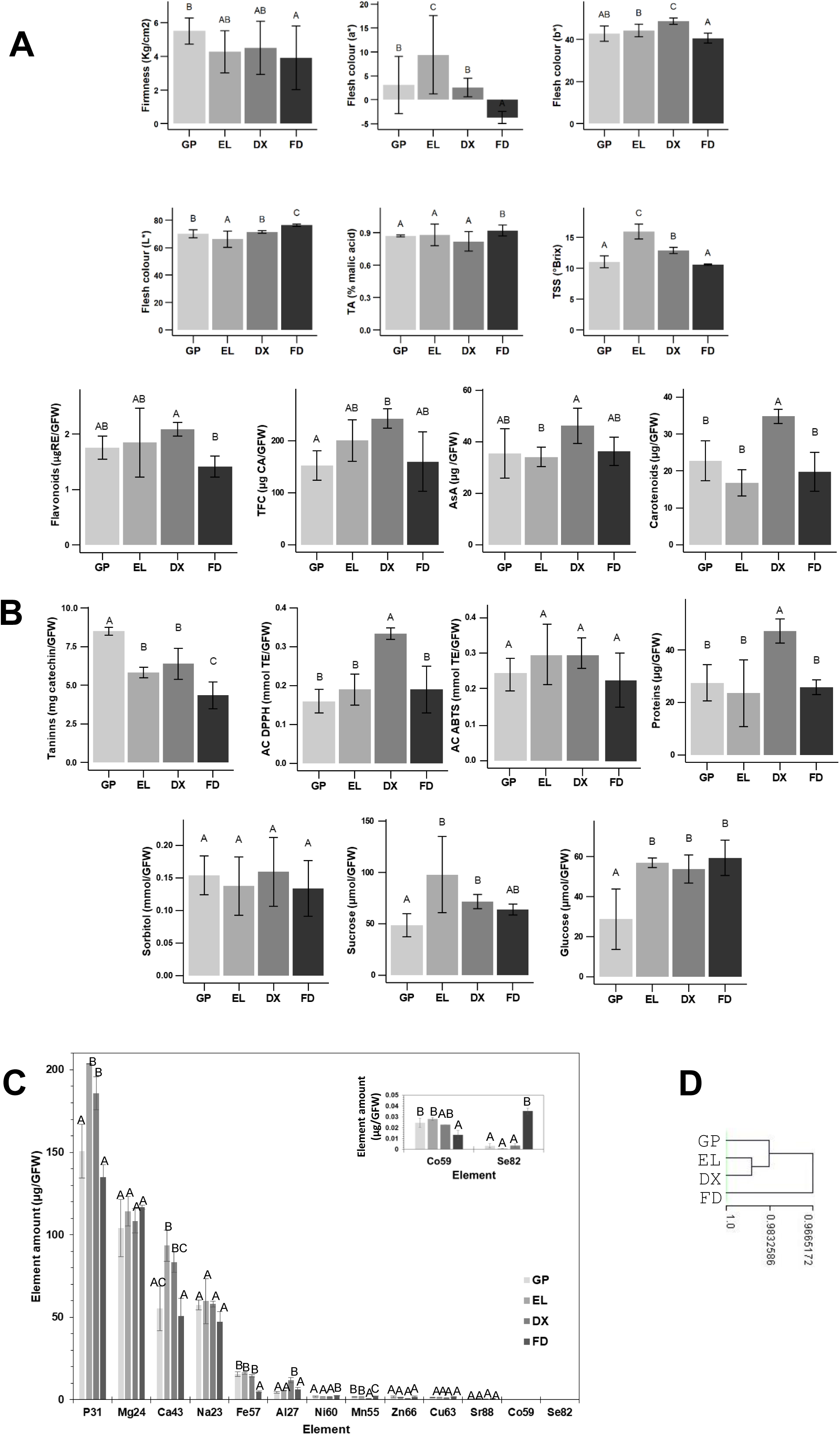
Biochemical and organoleptic characterization of four peach cvs grown and commercialized in Argentina. Fresh flesh was analyzed. Gold Prince (GP), Elegant Lady (EL), Dixiland (DX) and Flordaking (FD). **A.** Organoleptic characteristics **B**. Bioactive compound contents, antioxidant capacity (AC) and nutritional properties. **C.** Mineral composition of digested and filtered peach slices were assessed by inductively coupled plasma-mass spectrometry (ICP-MS). D. Hierarchical clustering analysis (HCA) of data collected from fresh fruits. Each content in B and C is expressed in a gram of fresh weight (GFW) basis. Each value is expressed as mean of at least five values ± standard deviation. Bars containing the same letters are not statistically significant different (ANOVA, Tukey test, *p ≤* 0.05). AsA, Ascorbic acid; CA, chlorogenic acid equivalent; RE, rutin equivalent; TA, titratable acidity; TE, trolox equivalent; TSS, total soluble solids.

Biochemical characterization was also conducted on fresh (F) peach fruits. The bioactive compound content was assessed by measuring total flavonoids, total tannins and ascorbic acid (AsA), total phenolic content (TPC) and carotenoids. Flavonoids exhibited a similar profile as TPC. While carotenoid content was the highest in DX, AsA levels in DX were only significantly different from those in EL. The highest amount of carotenoids in DX agrees with the highest b* parameter in the CIELab system evaluation. Tannins ranged between 8.50 ± 0.26 mg catechin/GFW in GP and 4.34 ± 0.88 mg catechin/GFW, in FD (Fig. 1B). The mid-season cvs showed the same tannin content. The antioxidant capacity (AC) of fresh fruits was evaluated using the DPPH and ABTS methods. The ABTS free radical scavenging activity did not depend on the cultivar, while the DPPH method revealed differences, with DX showing highest AC (Fig. 1B). Differences in results obtained by the ABTS and DPPH assays may be explained considering that ABTS is capable of reacting with both lipophilic and hydrophilic antioxidants, while DPPH can only be used in organic medium. With respect to nutritional properties, DX exhibited the highest amount of soluble proteins (47.27 ± 4.61 µmol/GFW, Fig. 1B). Regarding sugars and alcohol sugars, sorbitol levels were the same among cultivars, while sucrose and glucose differed. In agreement with the highest TSS content, EL showed the highest sucrose levels. It is noteworthy that sucrose molecular weight doubles that of glucose; therefore, sucrose contributes the most to the TSS content. EL, FD and DX exhibited the highest amounts of glucose (Fig. 1B).

The profiling of mineral elements of physiological importance was conducted in peach fruit. Among the major constituents of peach, P, Mg, Ca and Na were identified. K was also identified but it was not quantified here. The abundance of micronutrients, from highest to lowest, was a follows: Fe, Al, Ni, Mn, Zn, Cu, Sr, Co and Se. Differences in the amount of P, Ca, Fe, Al, Ni, Mn, Co and Se were found among cvs as shown in Fig. 1C. Peach nutritional properties not only are associated with the presence of vitamins, fiber and phytochemicals, but also to minerals. Mineral elements have essential roles in plant and human processes. As previously described, K is the most abundant mineral nutrient. Its concentration is greater than that of other fruits such as pear, apple and grapes (Kuang et al., 2022). Selenium is considered as essential microelement, found as selenoproteins and Se-amino acids such as seleno-methionine (SeMet) and seleno-cysteine (SeCys2) a, and it was present at very low levels in all cvs analyzed. Although mineral composition has been described for different cultivars of peach fruit (Iordanescu, et al., 2015; Manzoor et al., 2012), the presence of Al has only been described by da Rosa Louzada et al. (2022), Mitic et al. (2018) and in the current work. Ionome profiling is influenced by the soil and climate conditions (Debbarma et al., 2021). Since, peaches came from the same orchard, differences in the profiling may be consequence of differences in the genetics and period of growth.

Peach methanolic extract, dried and resuspended in DMSO were used to test the inhibition of enzymatic activities. Extracts from GP, EL, DX and FD could inhibit the activity of AMG, protease and AMY with a hyperbolic response (Table 1). IC50 was also estimated and the values are shown in Table 1. DX and FD extracts exhibited the greater percentage of maximum inhibition (Imax) for AMG, while EL and DX showed the highest AMY Imax. Conversely, GP presented the highest Imax for protease (Table 1). The amounts of DMSO extracts which give minimum and maximum IC50 values are equivalent to 11 and 44 g of fresh fruits. Proteases are enzymes that play a role in various chronic conditions, including inflammation and metastatic cancer. Synthetic inhibitors and polyphenols (flavonols, flavones, catechins and proanthocyanidins) have been used to inhibit these enzymes (Sartor et al., 2002). Imax described for the cvs analyzed agree with that of Chatos variety, which showed up to a 80% of Protease Imax (Belhadj et al., 2016). It has been reported that carbohydrate digestibility is associated with increased postprandial blood glucose levels. An approach to reduce postprandial hyperglycemia; and thus to prevent diabetes or manage it, is to inhibit the activity of carbohydrate-digesting enzymes such as AMY and AMG (Tundis et al., 2010). AMY Imax values are in line with results obtained for Glady’s variety (78.54 ± 2.75%) and for Chatos cv (95.83 ± 1.37%) (Belhadj et al., 2016). In contrast, other cvs did not detect AMY inhibition in the cvs tested (Mihaylova et al., 2021). Kandra et al. (2004) showed the proteases are inhibited by condensed tannins. Here, GP showed the smallest IC50 among cultivars and exhibited the highest tannin contents (Fig. 1B). Regarding AMG, differences in the inhibition were observed between cvs. These results agree with previous reports that shows the enzymeś inhibitor potential of peach is intensely dependent on the cultivar (Nowicka et al., 2018).

**Table 1.**
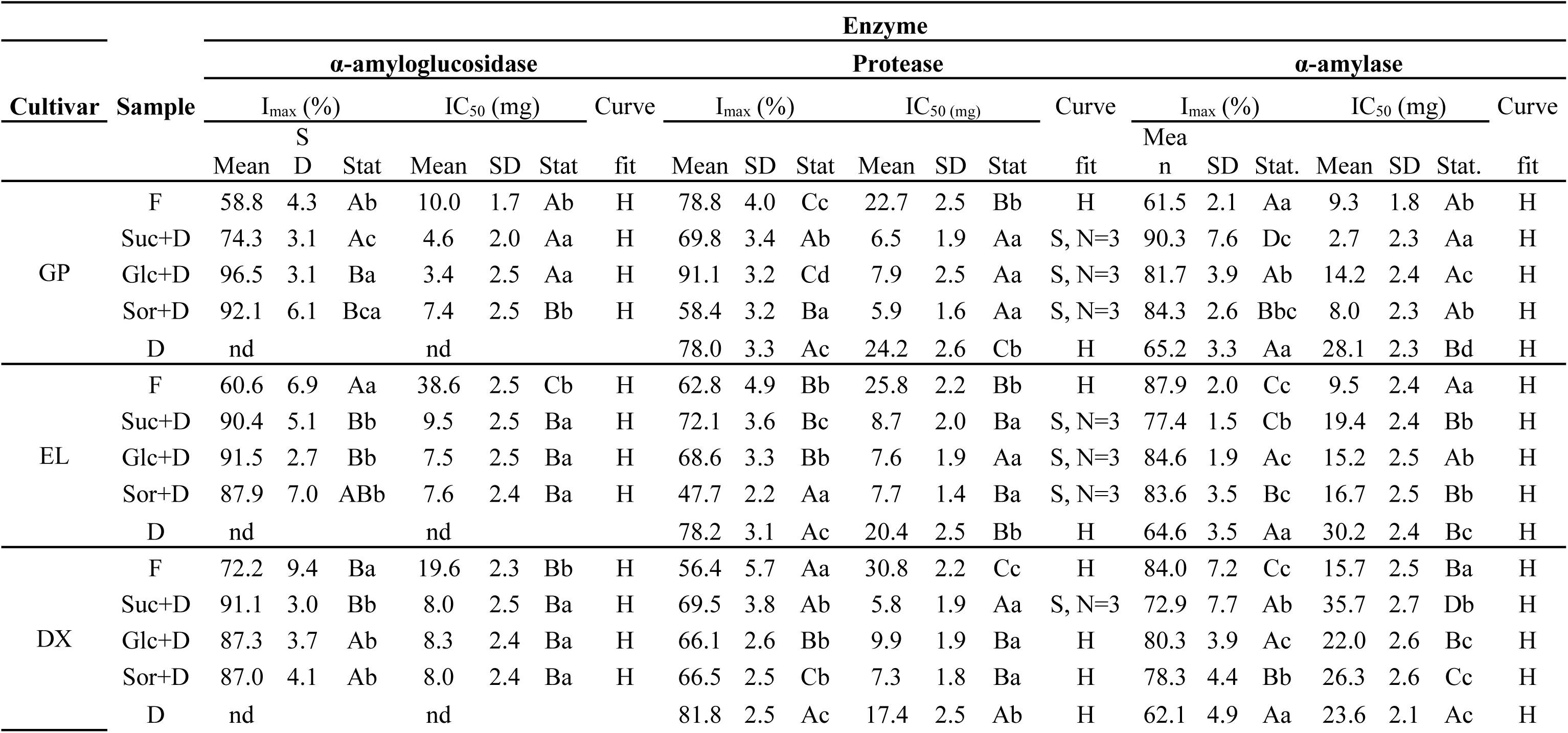
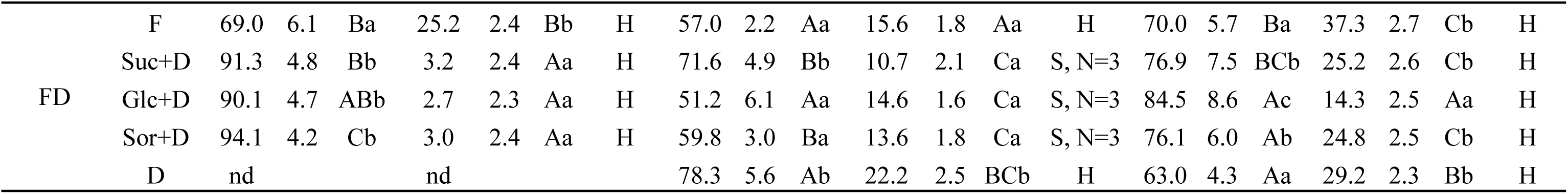
Enzyme inhibitions tests. α–amyloglucosidase (AMG), protease and α-amylase (AMY) inhibition power of DMSO extracts from GP, EL DX and FD. Extracts from fresh (F), dried (D) and OD treated previous hot air drying using sucrose (Suc+D), glucose (Glc+D) or sorbitol (Sor+D) peach slices were tested. For each enzyme, inhibition curves (percentage of inhibition vs extract amount) were fitted to sigmoidal (S) or hyperbolic (H) equations using SigmaPlot and used to estimate the IC50 and the maximum inhibition (Imax) of each extract. In the case of a sigmoidal curves, the Hill number (N) is provided. nd, not determined. Stat. shows the statistical analysis among samples (ANOVA, Tukey test, *p ≤* 0.05). Capital letters represent the statistical analysis performed to compare each type of sample (i.e. fresh) of the different cultivars. Lower cases indicate comparison of the different samples within each cultivar.

Fresh peach DMSO extracts were also tested against bacteria and yeast. The extracts from the four cvs inhibited the growth of the bacteria strains *Acinetobacter baumannii, Bacillus cereus, Escherichia coli, Enterobacter faecalis, Staphylococcus aureus* and *Salmonella typhimurium,* with the exception of FD which did not inhibit *E. coli* growth. The growth of *Candida albicans* and *C. tropicalis* was inhibited by all the extracts, whereas *C. glabrata, C. krusei* and *C. parapsilosis* were resistant to the extracts tested. The extracts from EL showed the largest inhibition zones against the pathogens tested, with the exception of *S. typhimurium.* In contrast, the extracts from FD showed the smallest inhibition. GP and DX presented an intermediate inhibitory activity compared to FD and EL (Table 2). Antimicrobial activity of peach fruit against *E. coli, S. aureus, B. cereus* and *C. albicans* agrees with those previously described for fruits from other peach cvs (Belhadj et al., 2016). Plant derived flavonoids have exhibited inhibitory activity against Gram-negative (*E. coli* and *A. baumannii*), Gram-positive (*Staphylococcus aureus* and *Enterococcus faecalis,*) and *Candida albicans* (Orhan et al., 2010), which could partially explain the inhibitions observed in this study. To the best of our knowledge this is the first time that inhibition of *C. tropicalis* is reported for peach flesh. *C. tropicalis* is an opportunistic fungal pathogen, a robust biofilm producer and many species are resistant to available antifungals. After *C. albicans, C. tropicalis* is the most important species responsible for candidiasis (Zuza-Alves et al., 2017).

**Table 2.**
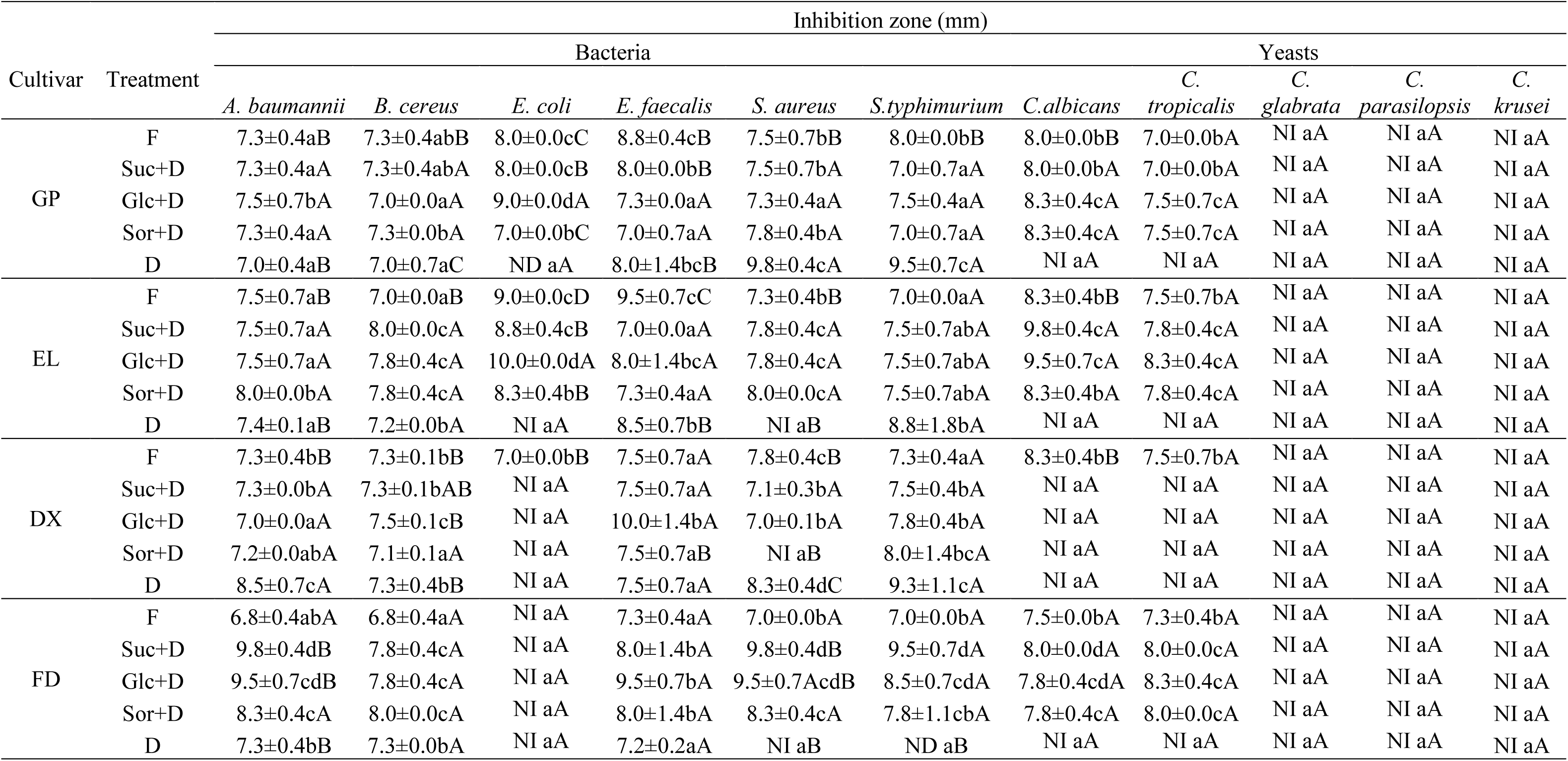
Antimicrobial activity of extracts from GP, EL DX and FD peach. Extracts were prepared using fresh (F), hot air dried (D) or OD treated slices followed by hot air drying using sucrose (Suc+D), glucose (Glc+D) or sorbitol (Sor+D) peach. The inhibition zones are expressed in millimeters as the mean of five replicates ± SD. For each strain and sample, values with at least one same capital letter are not statistical significant different among cultivars (ANOVA, Tukey test, *p ≤* 0.05). Within each cultivar statistical analysis is shown in lower cases. NI, no inhibition observed.

| Cultivar | Treatment | Inhibition zone (mm) |  |  |  |  |  |  |  |  |  |  |
| --- | --- | --- | --- | --- | --- | --- | --- | --- | --- | --- | --- | --- |
|  |  | Bacteria |  |  |  |  |  |  | Yeasts |  |  |  |
|  |  | *A. baumannii* | *B. cereus* | *E. coli* | *E. faecalis* | *S. aureus* | *S.typhimurium* | *C.albicans* | *C. tropicalis* | *C. glabrata* | *C. parasilopsis* | *C. krusei* |
| GP | F | 7.3±0.4aB | 7.3±0.4abB | 8.0±0.0cC | 8.8±0.4cB | 7.5±0.7bB | 8.0±0.0bB | 8.0±0.0bB | 7.0±0.0bA | NI aA | NI aA | NI aA |
|  | Suc+D | 7.3±0.4aA | 7.3±0.4abA | 8.0±0.0cB | 8.0±0.0bB | 7.5±0.7bA | 7.0±0.7aA | 8.0±0.0bA | 7.0±0.0bA | NI aA | NI aA | NI aA |
|  | Glc+D | 7.5±0.7bA | 7.0±0.0aA | 9.0±0.0dA | 7.3±0.0aA | 7.3±0.4aA | 7.5±0.4aA | 8.3±0.4cA | 7.5±0.7cA | NI aA | NI aA | NI aA |
|  | Sor+D | 7.3±0.4aA | 7.3±0.0bA | 7.0±0.0bC | 7.0±0.7aA | 7.8±0.4bA | 7.0±0.7aA | 8.3±0.4cA | 7.5±0.7cA | NI aA | NI aA | NI aA |
|  | D | 7.0±0.4aB | 7.0±0.7aC | ND aA | 8.0±1.4bcB | 9.8±0.4cA | 9.5±0.7cA | NI aA | NI aA | NI aA | NI aA | NI aA |
| EL | F | 7.5±0.7aB | 7.0±0.0aB | 9.0±0.0cD | 9.5±0.7cC | 7.3±0.4bB | 7.0±0.0aA | 8.3±0.4bB | 7.5±0.7bA | NI aA | NI aA | NI aA |
|  | Suc+D | 7.5±0.7aA | 8.0±0.0cA | 8.8±0.4cB | 7.0±0.0aA | 7.8±0.4cA | 7.5±0.7abA | 9.8±0.4cA | 7.8±0.4cA | NI aA | NI aA | NI aA |
|  | Glc+D | 7.5±0.7aA | 7.8±0.4cA | 10.0±0.0dA | 8.0±1.4bcA | 7.8±0.4cA | 7.5±0.7abA | 9.5±0.7cA | 8.3±0.4cA | NI aA | NI aA | NI aA |
|  | Sor+D | 8.0±0.0bA | 7.8±0.4cA | 8.3±0.4bB | 7.3±0.4aA | 8.0±0.0cA | 7.5±0.7abA | 8.3±0.4bA | 7.8±0.4cA | NI aA | NI aA | NI aA |
|  | D | 7.4±0.1aB | 7.2±0.0bA | NI aA | 8.5±0.7bB | NI aB | 8.8±1.8bA | NI aA | NI aA | NI aA | NI aA | NI aA |
| DX | F | 7.3±0.4bB | 7.3±0.1bB | 7.0±0.0bB | 7.5±0.7aA | 7.8±0.4cB | 7.3±0.4aA | 8.3±0.4bB | 7.5±0.7bA | NI aA | NI aA | NI aA |
|  | Suc+D | 7.3±0.0bA | 7.3±0.1bAB | NI aA | 7.5±0.7aA | 7.1±0.3bA | 7.5±0.4bA | NI aA | NI aA | NI aA | NI aA | NI aA |
|  | Glc+D | 7.0±0.0aA | 7.5±0.1cB | NI aA | 10.0±1.4bA | 7.0±0.1bA | 7.8±0.4bA | NI aA | NI aA | NI aA | NI aA | NI aA |
|  | Sor+D | 7.2±0.0abA | 7.1±0.1aA | NI aA | 7.5±0.7aB | NI aB | 8.0±1.4bcA | NI aA | NI aA | NI aA | NI aA | NI aA |
|  | D | 8.5±0.7cA | 7.3±0.4bB | NI aA | 7.5±0.7aA | 8.3±0.4dC | 9.3±1.1cA | NI aA | NI aA | NI aA | NI aA | NI aA |
| FD | F | 6.8±0.4abA | 6.8±0.4aA | NI aA | 7.3±0.4aA | 7.0±0.0bA | 7.0±0.0bA | 7.5±0.0bA | 7.3±0.4bA | NI aA | NI aA | NI aA |
|  | Suc+D | 9.8±0.4dB | 7.8±0.4cA | NI aA | 8.0±1.4bA | 9.8±0.4dB | 9.5±0.7dA | 8.0±0.0dA | 8.0±0.0cA | NI aA | NI aA | NI aA |
|  | Glc+D | 9.5±0.7cdB | 7.8±0.4cA | NI aA | 9.5±0.7bA | 9.5±0.7AcdB | 8.5±0.7cdA | 7.8±0.4cdA | 8.3±0.4cA | NI aA | NI aA | NI aA |
|  | Sor+D | 8.3±0.4cA | 8.0±0.0cA | NI aA | 8.0±1.4bA | 8.3±0.4cA | 7.8±1.1cbA | 7.8±0.4cA | 8.0±0.0cA | NI aA | NI aA | NI aA |
|  | D | 7.3±0.4bB | 7.3±0.0bA | NI aA | 7.2±0.2aA | NI aB | ND aB | NI aA | NI aA | NI aA | NI aA | NI aA |

C. tropicalis is more phylogenetically closely to C. albicans than C. glabrata, C. krusei and *C. parasilopsis*; explaining the common susceptibility of *C. albicans* and *C. tropicalis* to the extract tested. Additionally, while inhibition of the infective endocarditis bacteria *Enterococcus faecalis* has been reported for extracts from peach bark (Aziz & Rahman, 2013) and leaves (Koyu et al., 2020), inhibition of the edible flesh has not been previously described. Moreover, the inhibition against *A. baumannii* shown by GP, EL, DX and FD is of great relevance. This opportunistic bacterium is a major concern in the hospital environment due to its multidrug resistance. Antibacterial capacity has been demonstrated for seed oils from *P. armeniaca* and *P. avium* but not from *P. persica* (Fratiani et al., 2021). Finally, HCL tree was created using all the parameters measured for fresh fruits (Fig. 1D). EL and DX, mid-season peaches belonged to the same branch of the tree, while GP and FD were more distant. The four cvs analyzed were originated in the USA (Okie, 1998) and were cultivated in the same orchard; thus, clustering obtained reflects the impact of the pedigree and the weather on the organoleptic and nutraceutical characteristics of the cvs.

### 3.2. Osmotic Dehydration of peach fruit

#### 3.2.1. Moisture content, water loss, solute gain and colour are dependent on peach cultivar

GP, EL, DX and FD peach fruit were subjected to OD treatment using three different solutions containing 46-47°Brix Glc, Suc or Sor for 3 h at 40 °C in the presence of 2% (w/v) CaCl2 followed by D. Slices obtained were compared with those of F and D fruit. Representative images are shown in Fig. 2 for each cv. Peach fruit are characterized by high moisture content (MC, 86-89%); OD rendered an average MC of 56% for all cvs (Fig. 3A). Afterwards, the slices were dried until a 13% MC. In parallel, fruits were exclusively hot-air dehydrated for a longer period of time to reach a 14% MC. While differences in the MC of the different cvs were observed before and after the treatments, statistical analysis revealed that within each cv, the same MC is obtained within OD fruits independently the HS used, which finally renders OD+D slices with the same MC (Fig. 3A, Box). Table 3 presents the analysis of the effect of cv and treatment in each parameter measured in this work.

**Fig. 2.**
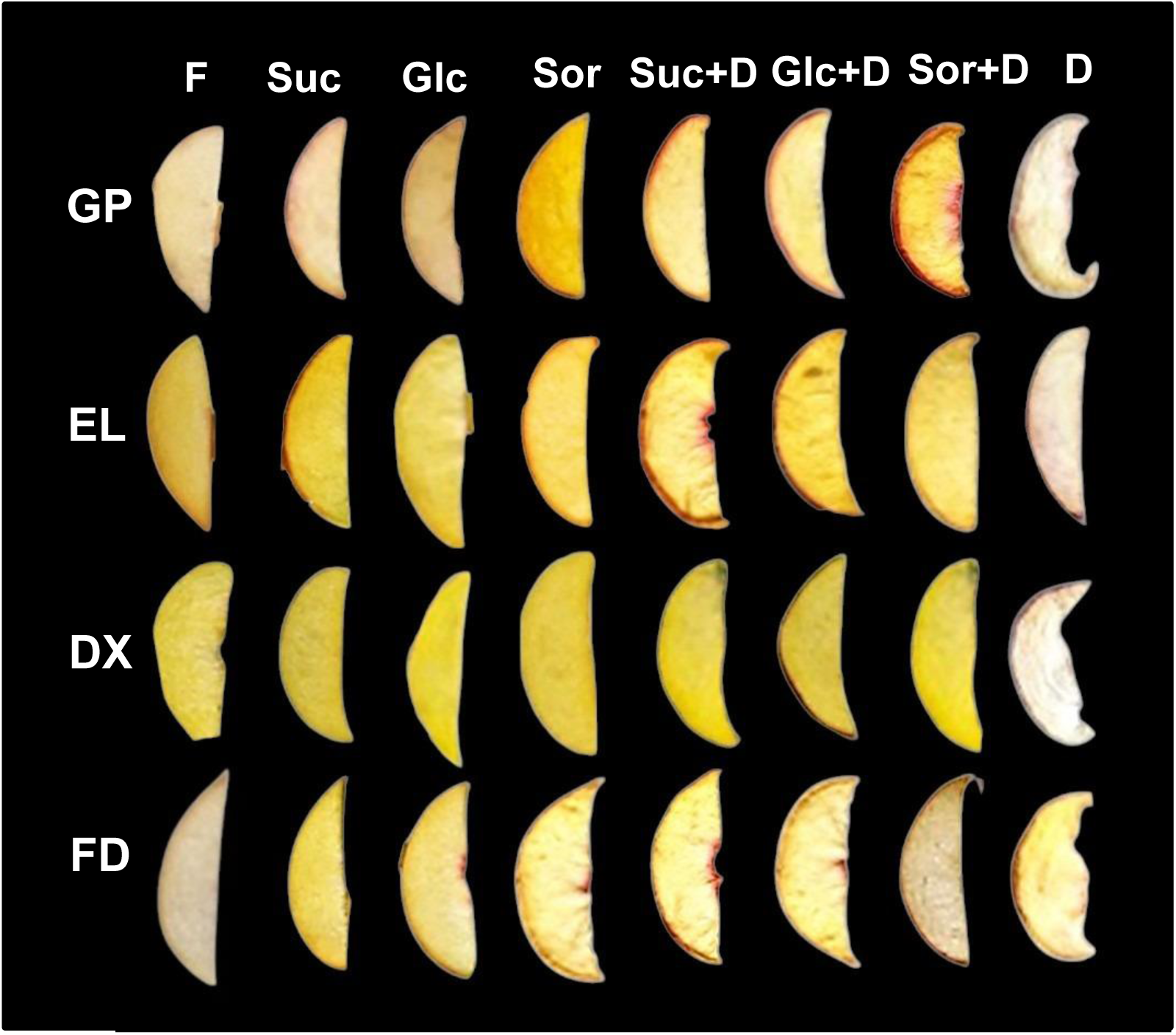
Representative images of peach slices analyzed. Gold Prince (GP), Elegant Lady (EL), Dixiland (DX) and Flordaking (FD) cultivars were used and analyzed as recently harvest (fresh, F) or after being incubated in HSs containing sucrose, glucose or sorbitol (Suc, Glc and Sor), followed by conventional heat air drying (Suc+D, Glc+D and Sor+D) or after being exposed to conventional air drying (D).

**Fig. 3.**
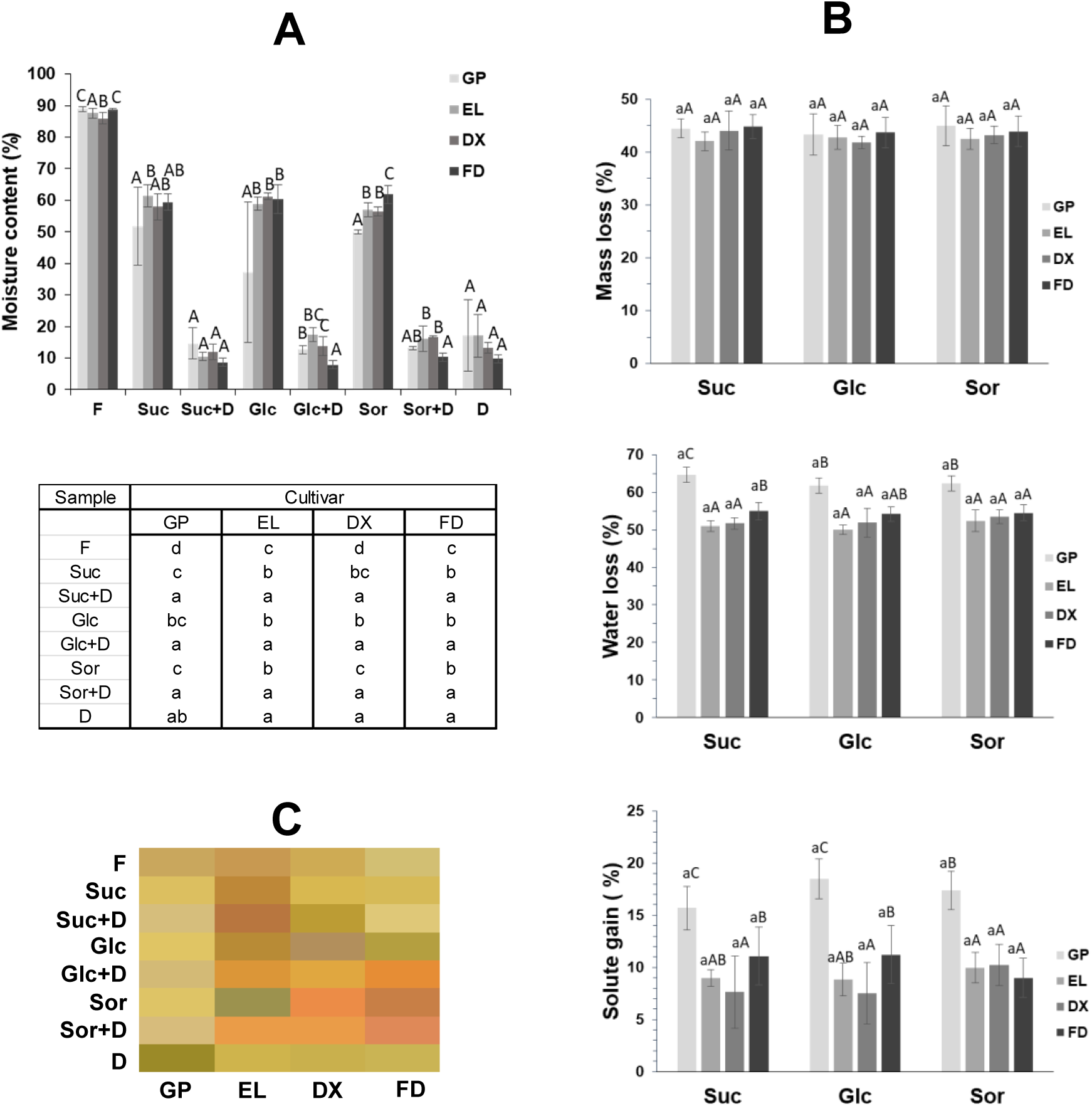
**A.** Moisture content of fresh (F), osmotic dehydrated with Suc, Glc or Sor (Suc, Glc and Sor), followed by conventional heat air drying (Suc+D, Glc+D and Sor+D) and conventional air dried (D) peach slices of the following cultivars Gold Prince (GP), Elegant Lady (EL), Dixiland (DX) and Flordaking (FD). Bars represent the mean *±* standard deviation of at least five replicates. Among each type of sample, bars with different letters indicate statistically significant differences among cultivars (ANOVA, Tukey test, *p ≤* 0.05). The table below the graph shows the statistical analysis comparing samples within each cultivar (ANOVA, Tukey test, *p ≤* 0.05). **B**. Mass transfer parameters of peach slices subjected to osmotic dehydration; mass loss, water loss and mass gain (expressed in a fresh weight basis). Bars represent the mean *±* standard deviation of at least five replicates. Bars with at least one same letter are not statistically significant different (ANOVA, Tukey test, *p ≤* 0.05). For each OD treatment, capital letters reveal differences among cultivars. Small letters indicate comparisons among dehydration treatments with sucrose, glucose or sorbitol within each cultivar. **C.** Color map of representative (media of measurements) La*b* parameters. La*b* parameters were mapped to colors on a figure using the “Colorspace” package from “Rstudio”.

**Table 3.**
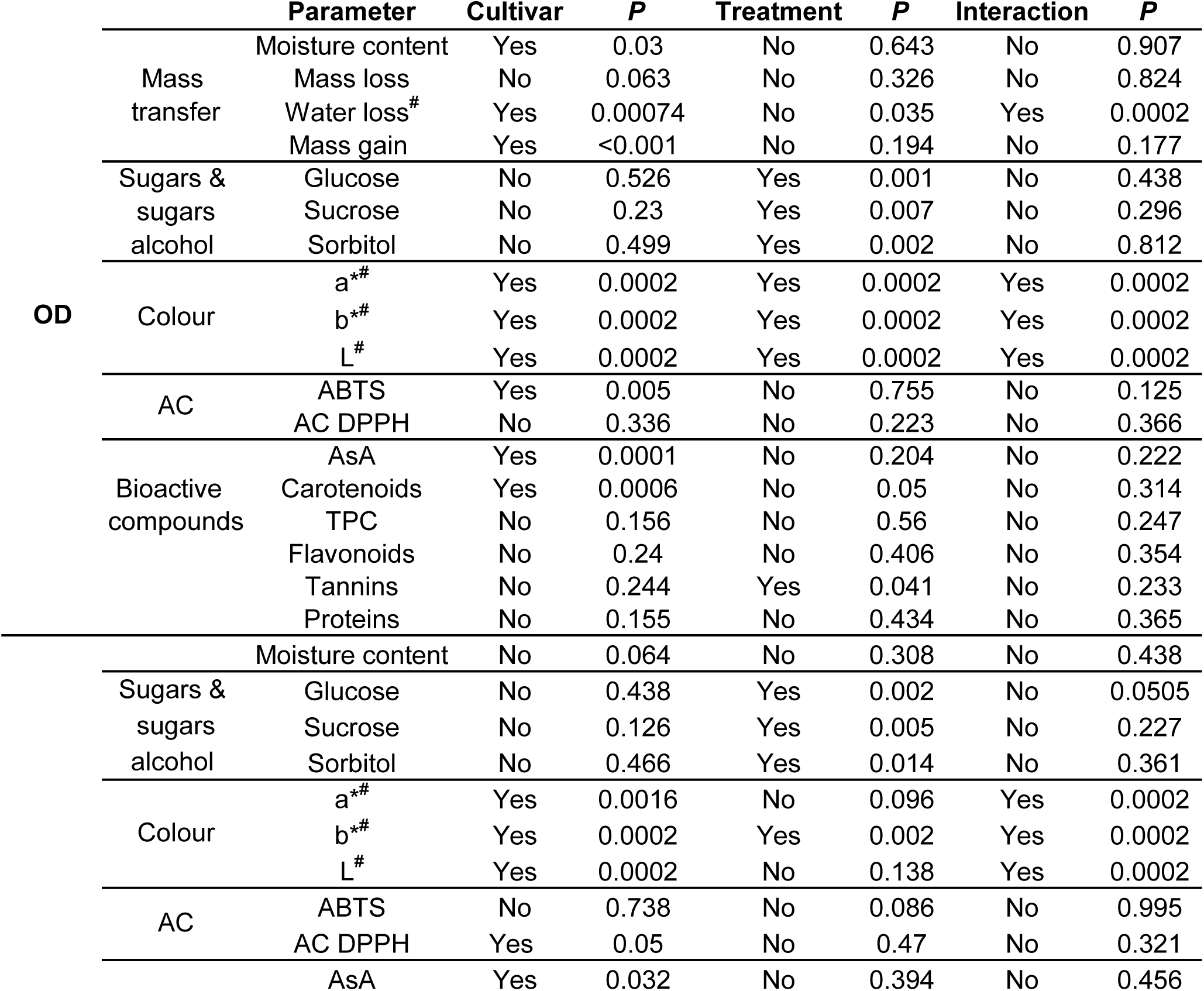
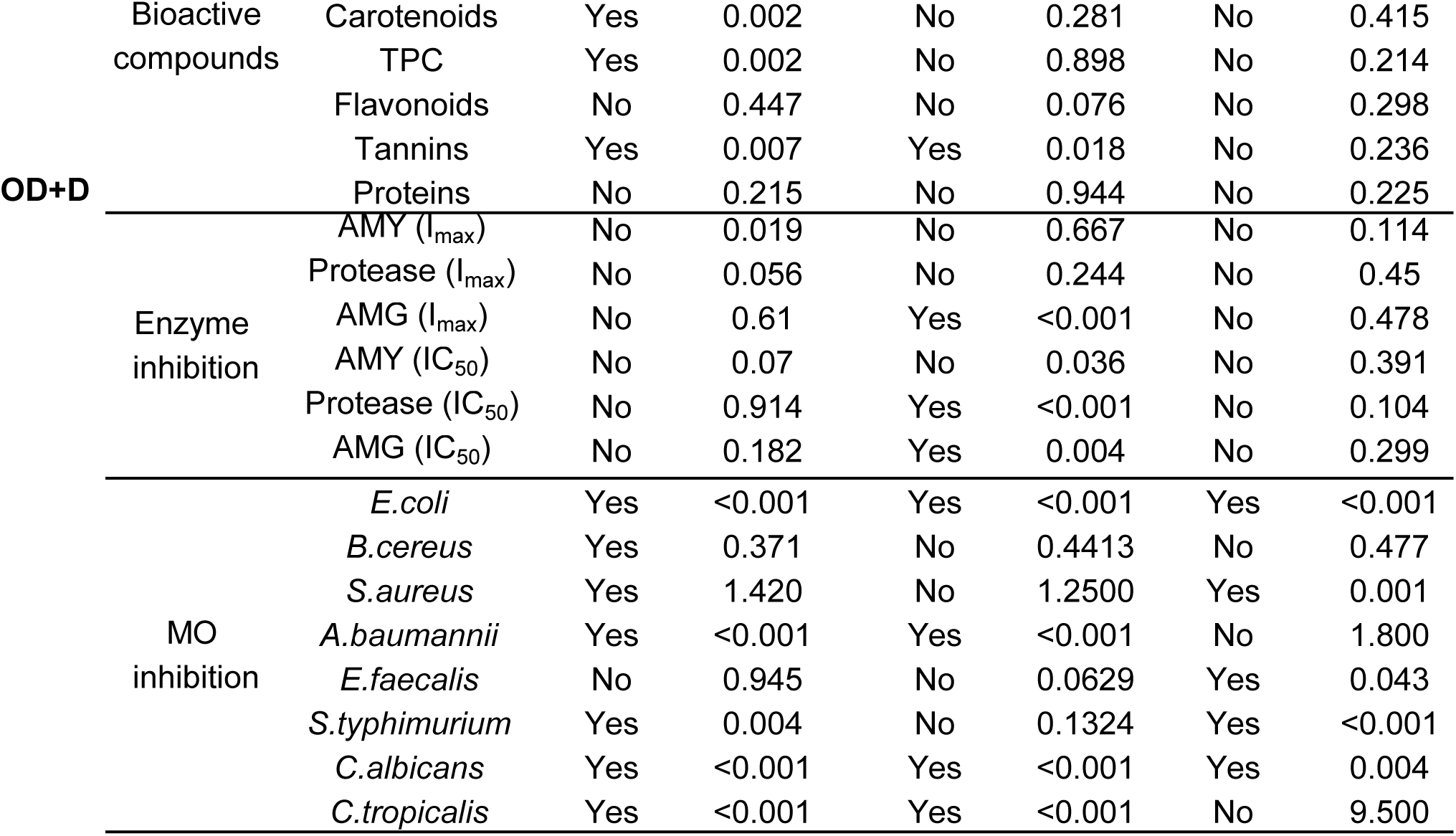
Analysis of the effect of cultivar (GP, EL, DX and FD) and OD-D or OD-D treatments in each parameter measured in the current work. Two way ANOVA was conducted (*p ≤* 0.05). Cases in which equal variance test failed are marked with # and data was analyzed using permutational multivariate analysis of variance (PERMANOVA), *p ≤* 0.05.

Mass Loss (ML), water loss (WL) and solute gain (SG) were computed to characterize the OD process (Figure 3B). The ML was not affected neither by the cv nor the HS. WL ranged between 50.0±1.3% for EL using Glc and 64.6±2.0 for GP using Suc. SG ranged between 7.5±3.0% in DX and 18.5±1.9% in GP, both dehydrated with Glc solutions. Within each cv tested, WL and SG did not exhibit variations with OD solution used. However, different results were obtained depending on the cv (Table 3); GP exhibited the highest WL and SG, followed by FD (Fig. 3B). Using a 10-mm-thick peach slices, a 1:5 fruit:solution ratio, and sucrose 30 °Brix solution during 3h at 25°C, Wang et al. (2023) achieved a SG of 7.7% and a WL of 17.3 %. After 5h of incubation a SG of 9.5 (%) was obtained; and using isomaltooligosaccharide a 23.2% was achieved after 5h. On the other hand, for 5-mm-thick slices, 1:5 fruit:solution ration, 50 °Brix sucrose, for 3h at 35°C Singh Yadav et al. (2012) obtained a SG of 8.3% and a 19.4% WL. The maximum WL of 31.3% was achieved using 70 °Brix sucrose for 4 h at 55°C. Our results agree with these reports in the SG, while the WL obtained here is higher. Nevertheless, Giangiecomo et al. (1987), described a 45 % WL using corn syrup of °70 Brix and a 1:5 fruit:solution ration, which is closer to our values. The OD in the current work was conducted in the presence of calcium and 1:10 fruit:solution ratio. The presence of calcium salts in the OD solution rises the WL and slightly decreases the SG. Calcium is retained in the fruit cell walls altering their structure which increases tortuosity and local viscosity; and then, decreases the up-take of sugars of the food. Calcium also improves the texture of the product (Espinoza Estaba et al., 2006). Additionally, increase in fruit:solution ratio may also contributes to in the augmentation WL.

Colour parameters of peach F and treated slices were recorded and shown in Supplementary Table 1. Color map of representative La*b* parameters is shown in Fig. 3C. Changes in colouration are both dependent on the cv and OD solution used (Table 3). Regarding yellowness (b*), OD+D resulted in higher values than F in EL (Glc+D, Sor+D), DX (Suc+D, Glc+D, Sor+D) and FD (Glc+D). b* also increased in D from EL, FD and GP. Increases redness (a*) with respect to F were observed in Glc+D and Sor+D in DX and FD. In contrast a* decreased in D (DX, GP). In GP, L (lightness) significantly decreased in D but increased with all OD+D treatments. Nevertheless, the other cvs exhibited a contrasting behavior. The ΔE*76 parameter, which represents the differences in the total colouration perceived by the human eye was calculated for each treated sample in comparison with F (Supplementary Table 2). The colour of treated slices changed with respect to the F and for all cases changes were classified has highly noticed by the human eye. While in GP and EL there was a slight variation among OD solutions used, DX and FD were more variable. In the case of DX and FD, greater changes were perceived when treated with Sor, and the smallest perception was noticeable when Suc is used. Colour is a complex character influenced by different factors such as pigment composition, oxidative stability, sugar interactions, and structural modifications (Chauhan et al., 2011; Cortez et al., 2016). Colour analysis and images indicates that browning reactions are limited in most treatments, with some HSs playing a protective role in colour retention. Both cv and HS show interaction in final appearance of the dehydrated peach slices. Future sensory studies could help to elucidate consumers’ preferences on dehydrated peach.

#### 3.2.2. A higher metabolites and AC retention is observed for OD+D peach slices with respect to D peaches

To assess the quality of the OD+D peach slices, antioxidant, bioactive and nutritional properties were evaluated. As expected due to a decrease in the water content of the samples (Fig. 3A), increases in all the parameters tested in OD, OD+D and D slices were observed when results were expressed in a FW basis (Supplementary Figs. 2-5). Therefore, to identify the changes in fruit composition and capabilities due to OD and D process, values were calculated in a DW basis. Given that in OD and OD+D a significant SG is produced (Fig. 3B), a correction factor considering the increase in DW due to Suc, Glc or Sor up-take into the slices was applied to represent the net variation in the metabolite or capacity tested. Only when the metabolite evaluated was the one used for OD, the correction was not applied (i.e. Glucose in Glc+D and Glc samples, Fig. 4). Net Suc up-take into the fruits were confirmed in Suc and Suc+D slices of all cvs. The same behavior was observed for Glc and Sor in slices incubated with these osmolytes. In agreement, the content of Glc, Suc and Sor were influenced by the treatments in OD and OD+D fruits (Table 3). In contrast, these parameters were not influenced by the cv. Metabolic interconversion between Suc and Glc is observed in treated fruits; in DX, FD and GP, the net content of Glc is increased with respect to F slices when Suc is used in the HS revealing Suc hydrolysis within the peach slice; and an increase in Suc is observed in DX when Glc is the HS, indicating up-take that part of the up-taken Glc is used for Suc synthesis. Regarding D, the net content of Sor was not affected in most of the D peaches (DX, EL and FD); however, Suc decreased in FD and EL, and Glc decreased in FD, probably due to enhanced respiration during drying in the heat (Fig. 4).

**Fig. 4.**
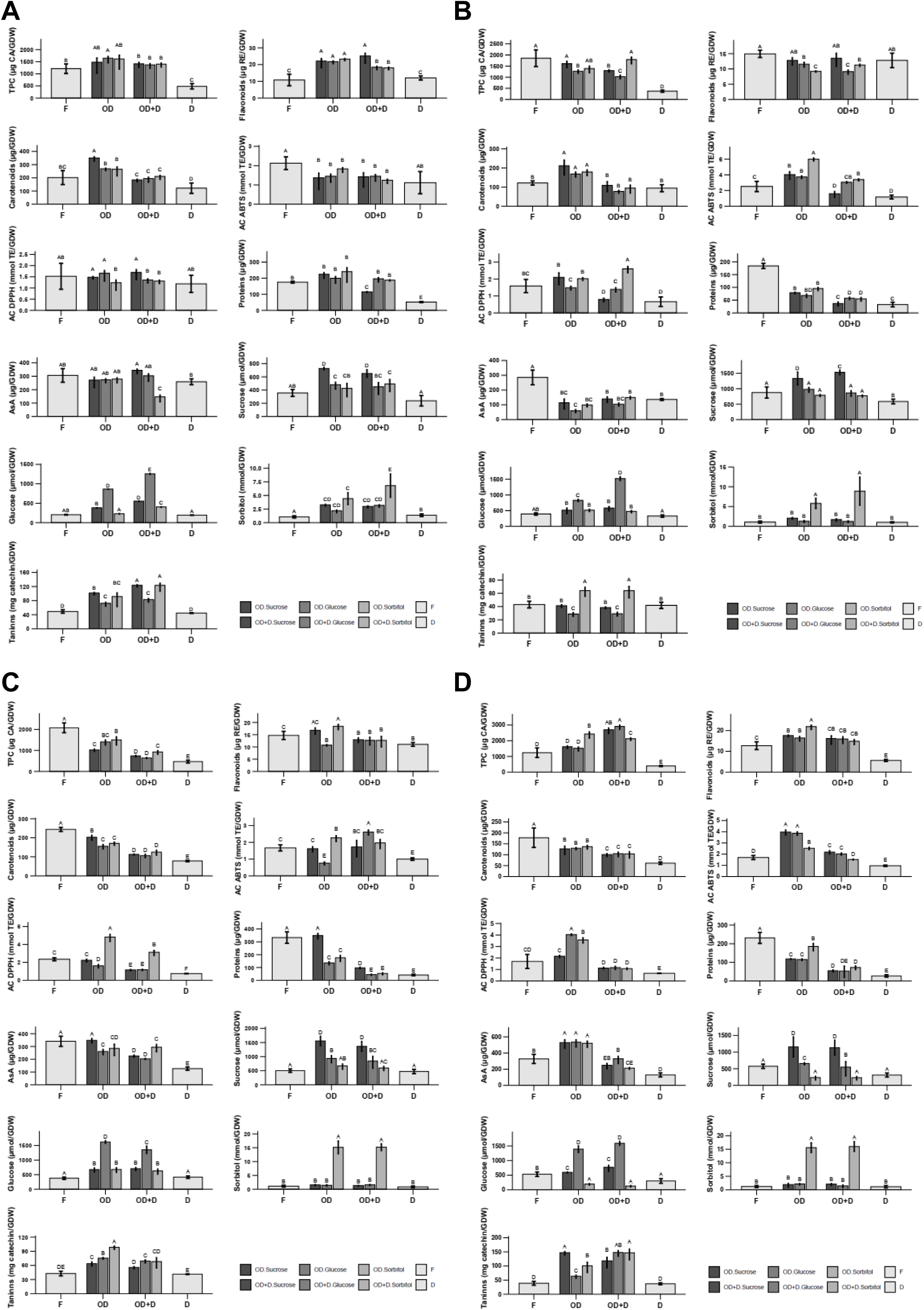
Bioactive compound contents, antioxidant capacity (AC) and nutritional properties of peach fruit dried (D), or incubated with HSs of Suc, Glc or Sor (OD) followed by hot air drying (OD+D). Bars with at least one same letter are not statistically significant different (ANOVA, Tukey test, *p ≤* 0.05). Results are expressed in a GDW basis. **A.** Gold Prince, **B.** Elegant Lady, **C.** Dixiland, **D.** Flordaking.

With respect to bioactive compound, with the exception of tannins, the behavior was not dependent of the type of HS in OD and OD+D slices (Table 3). Depending on the cv, TPC was decreased (DX, EL), increased (FD) or remained constant (GP, EL) with respect to F (Fig. 4.). Despite some diminishes, the amounts were always greater than in D, which showed significant decreases in all cvs with respect to F (up to a 0.20-fold in EL). Phenolic compounds are proposed as indicators of fruit quality since they contribute to the taste and visual appearance of the fruits. TPC are an important source of the AC in peach (Lara et al., 2020). Prolonged exposure to 58°C decreases the content of TPC in D, irrespectively of the cv. Thus, the increase in the heat intensity (including temperature and duration of exposure) conducts to enlarged thermal degradation of TPC (Nowicka et al., 2025). In contrast, OD has a variable impact on TPC. Although flavonoids followed a similar trend to TPC, differences in the contents indicate there are other phenolics such as chlorogenic acid mainly contributing to the TPC in these cvs (Fig. 4). In agreement, previous studies reveal that flavonoids are not the main TPC in peach fruit (Saidani et al., 2017). Peach cvs differ in the content and the TPC profiling. The diverse phenolics show variations in their temperature stability and also may differ in their diffusion coefficient (Nowicka et al., 2025).In this respect, it is expected variations in the responses among cvs.

In OD+D fruits Flavonoids were increased in GP and DX, invariable in FD, and decreased in DX and in EL with respect to F. Flavonoids diminished in D slices of DX and FD. In the case of GP and FD, flavonoid content was higher in OD+D than in D (Fig. 4). Flavonoids are a wide group of fifteen-C phenylpropanoid core derived from the shikimic acid pathway. The exhibit a wide range of activities, among which antioxidant, antibacterial, antifungal, antiviral, antitumor and anti-inflammatory can be mentioned (Eun & Maruf, 2017). Peach contain catechin, epicatechin, epigallocatechin, procyanidins, rutin, quercetin-3-glucoside, quercetin-3-galactoside and kaempferol-3-rutinoside (Lara et al., 2020). On one hand, sugars have been shown to induce the synthesis of flavonoids (Morkunas et al., 2005, Tsukaya et al, 1991, Lv et la., 2022). Therefore, the observed increases of flavonoids in GP, DX and FD in OD slices may be attributed to up-regulation of their biosynthesis. Afterwards, during air-drying in OD+D, these compounds are maintained or slightly decreased. Conversely, in EL, flavonoids are not induced in OD (Fig. 4). Thermal processes negatively or positively impact on the flavonoid content of a wide range of fruits and vegetables, as extensively reviewed by Eun & Maruf (2017). Our results display a wide type of responses to OD and to D, showing that an intricate combination of factors, which undoubtedly depend on the cv, are involved in the regulation of flavonoids levels in peach slices.

AsA did not depend on the type of osmolyte used but varied with cv, with diminishing or preserving behavior in OD+D with respect to F. In D, AsA decreased in FD, DX and EL with respect to F. When comparing OD+D with D, OD+D exhibited higher amounts of AsA in all cvs expect of EL. AsA is a heat sensitive molecule; thus, the higher drying temperature (58°C vs 40°C) and the longer period of heating of D was expected to conduct to a lower content of AsA in D with respect to OD+D. Besides a leaking of AsA to the HS has been proposed during the OD, followed by a degradation during the following hot-air-drying (Yadav and Singh, 2014); this is not the case for GP, FD and DX, in which AsA is not decreased in OD+D with respect to D (Fig. 4). Differences in fruit ultrastructure and composition among peach cvs may account for this variations in the AsA retention. In addition, variations in the type of fruit also conduct to different responses. While in apricot Sor had a better than Suc performance regarding AsA retention (Riva et al., 2005), the response in peach slices depends on the cv (Table 3).

Carotenoids did not vary in EL and GP in OD+D with respect to F, but decreased in DX and FD. The reduction in the carotenoids in OD+D ranged between 10.5 % for Suc+D in EL and 56.4 % for Glc+D in DX. In D slices decreases in carotenoids in GP, DX and FD were more severe than in OD+D (from 22.4 % in EL and 67.9 % in DX, Fig. 4). For reference, guava halves showed a 44% decrease in carotenoids when incubated with 60% Suc for 2h followed by hot-air-dried and a 75% without sugar pre-treatment (Sanjinez-Argandona et al., 2005). Thus, the OD process alleviates carotenoid degradation. It has been proposed that sugars in the surface of the foods create a barrier layer restraining the reaction between oxygen and carotenoids (Luchese et al., 2015; Tonon et al., 2007). While carotenoids retention in kiwifruit was greater using Sor than Suc (Bialik et al., 2020); carotenoid preservation here was dependent on the cv but not on the HS. Kroehnke et al. (2021) proposed that a higher solid gain could restrict the O2 entrance to the dried food. Our results are in agreement with this concept, since GP exhibited the highest SG and did not showed a decrease in the carotenoid (Figs. 3 and 4).

Tannins varied depending on the cv and the treatment (Table 3) in OD+D slices. The general trend was of increase in OD+D peaches with respect to F, while in D remained constant (Fig. 4). The increase in tannins in OD+D slices is a remarkably feature. Different stress conditions such as pathogens, UV-B light and wounding induce tannins biosynthesis (Mellway et al., 2009). Both anthocyanins and tannins share common biosynthetic precursors since they derive from the shikimic acid and phenylpropanoid pathways with many characterized transcription factors. Here, the content in tannins only occurs in slices exposed to HS; therefore, sugars might be involved in such an induction and not the heat. Sugars are molecules signalling probed to induce anthocyanins synthesis through WD40 and other transcription factors modulating the expression of key enzymes such as dihydroflavonol reductase and anthocyanidin reductase also involved in tannins biosynthesis (Das et al., 2012). Thus, the presence of sugars in the incubation solution stimulates tannins biosynthesis through sugar signaling processes. Moreover, the presence of calcium may also trigger other signaling mechanisms. While condensed tannins have originally been considered nutritionally undesirable and responsible for the astringency to the foods; their role in foods have been revised. Condensed tannins, as different phenolics and flavonoids, act as strong antioxidants *in vitro* and *in vivo* (Hagerman et al., 1998; Gourlay & Constabel, 2019). They also exhibit anti-inflammatory, antimicrobial, anti-diabetic and anti-obesity activities (Santos-Buelga & Scalbert, 2000). Increases of up-three fold in the amount of tannins have been observed after OD+D (Fig. 4) in FD and GP, rendering slices with 70.75± 6.41 mg/GPF (Supplementary Figure S5), with similar values to those observed in apples, plums and raspberries, a fifth of the amount of milk chocolate and a sixth part of that in dark chocolate, one of the richest sources of tannins (Arts et al., 2000).

The total soluble protein content was not dependent on the cv or HS. Decreases in these macromolecules were observed in all OD+D samples with respect to F. Nevertheless, in general, the levels were greater than in D slices (Fig. 4). The reduction in total soluble proteins was not dependent on the cultivar neither the HS. This reduction could be related to protein precipitation and denaturation which is higher in fruit exclusively exposed to hot-air-drying.

Finally, AC was evaluated by ABTS and DPPH assay. In D peaches, the AC decreased in all cvs. In OD+D slices, DPPH scavenging capacity depend on the genotype, with responses of increase, decrease or no variation with respect to F. When AC was tested by the ABTS method, a variety of responses were observed for many samples. Besides GP, for most of the samples the AC was higher in OD+D than in D slices (Fig. 4).

#### 3.2.3. Osmotic dehydration treatments followed by conventional heat-drying improves the inhibition towards protease, α-amylase and α-amyloglucosidase

The inhibition of different enzymes activities was assayed in OD+D and D extracts from GP, EL, DX and FD (Table 1). A greater Imax of AMG was achieved for OD+D with respect to F, irrespectively of the cv and HS, reaching almost complete inhibition (96.5±3.1% in Glc+D for GP). In line with these results, decreases in IC50 of AMG were observed in all cases with the exception of Sor+D, GP (Table 1). A similar trend was observed for AMY, which Imax was exacerbated in OD+D of GP and FD, and remained unaltered in EL and DX (Table 1). Therefore, OD+D improves the nutraceutical feature of the peaches. In the case of the inhibition of the protease activity, Imax was increased in DX with respect to F and using some HS in GP, EL and FD. Nevertheless, IC50 of protease was diminished for most of the cvs, revealing that a less amount of the food is needed to reach the complete inhibition (Table 1). Changes in the kinetics (from hyperbolic to sigmoidal) of the inhibition were observed for protease, revealing that probably more than one metabolite is contributing to the enzyme inhibition in these cases. The inhibition of enzymes involved in carbohydrate metabolism is a strategy to manage diseases related to hyperglycemia and obesity (Tundis et al., 2010). On the other hand, protease activity has been related to inflammatory process. The consumption of fresh fruits have been documented to play an important role in modulating metabolic risks factors (Sun & Miao, 2020). Here, in the current work it is shown that not only fresh peach inhibits AMG, AMY and protease activities but also in the osmotically-dehydration version of peach, with a better performance in most cases.

#### 3.2.4. Dehydration processes modify the antimicrobial capacity of peach slices

Changes in the inhibition zones of bacteria and yeasts growth in DMSO extract from peach were observed after OD+D and D (Table 2). Significant changes ranged in magnitude. Regarding yeasts, the inhibition depended on the cv. While in DX, a loss of the inhibition was observed when *C. albicans* and *C. tropicalis* were tested, increased inhibition with respect to F was observed in GP, EL, and FD. Conversely, D did not inhibit the growth of this yeasts. In the case of bacteria, the response was very variable depending on the cv. FD exhibited significant increases in the inhibition zones of *A. Baumannii, B. cereus, E. faecalis, S. aureus* and *S. typhimurium*. On the contrary, GP trend in OD+D was of decrease or unchanged; only improving the inhibition of *A. baumannii* and *E. coli* in Glc+D. EL and DX, presented intermediate responses, showing increases, decreases and no responses against the bacteria. In general, the behavior of extracts from D slices showed a trend in which only a in a third of the cases the inhibition activities against bacteria was improved, denoting that prolong heat treatment at 58°C has a negative effect in the inhibition power of the peaches.

#### 3.2.5. Mineral composition is altered in OD+D which are enriched in Ca

Ionomics of OD+D fruits was conducted by ICP-MS and compared to that of F fruit. Heat map showing the relative abundance of minerals in osmotic dehydrated peach slices (OD+D) with respect to fresh fruit is shown in Fig. 5. Mineral contents of each sample are expressed as µg/GDW and shown in Supplementary Table S1. Of the 132 pairs analyzed, 91 % showed changes in their relative amount, with all the elements analyzed exhibiting variations after the process. In general, and as expected, the relative amount of most of the minerals in OD+D decreased compared to F. This behavior can be explained by a net decline in the amount of the element, due to an efflux of the minerals dissolved in the water that exits the fruit during the OD procedure. On the other hand, part of this decrease in the concentration can be attributed to changes in the dry weight, resulting from the increase in the dry mass due to the uptake of the Suc, Glc or Sor. Conversely, despite this last statement, a net increase in Calcium was observed in all OD+D samples which ranged between 1789.08 ± 320.35 and 6306.79 ± 1132.22 µg/GDW and 1.5±0.26 and 5.5± 0.84 mg/GFW. The average increase in Calcium was of 6.8 fold and the maximum increase of 15.6-fold was recorded in Glc+D compared to GP. Thus, the presence of Calcium in the HS used for OD leads to an enrichment of this essential microelement in the dehydrated peach slices. For comparison, milk contains 1.25 mg/g of calcium (https://fdc.nal.usda.gov/food-details/173441/nutrients) and almonds from Spain have 1.9–6.7 mg/g (Yada et al., 2012). Therefore, the concentration of calcium in the OD+D peach slices is in the range of other foods rich in this element. While milk is consumed in higher amount, the consumption in grams of OD+D slices may be equivalent to those of almonds.

**Fig. 5.**
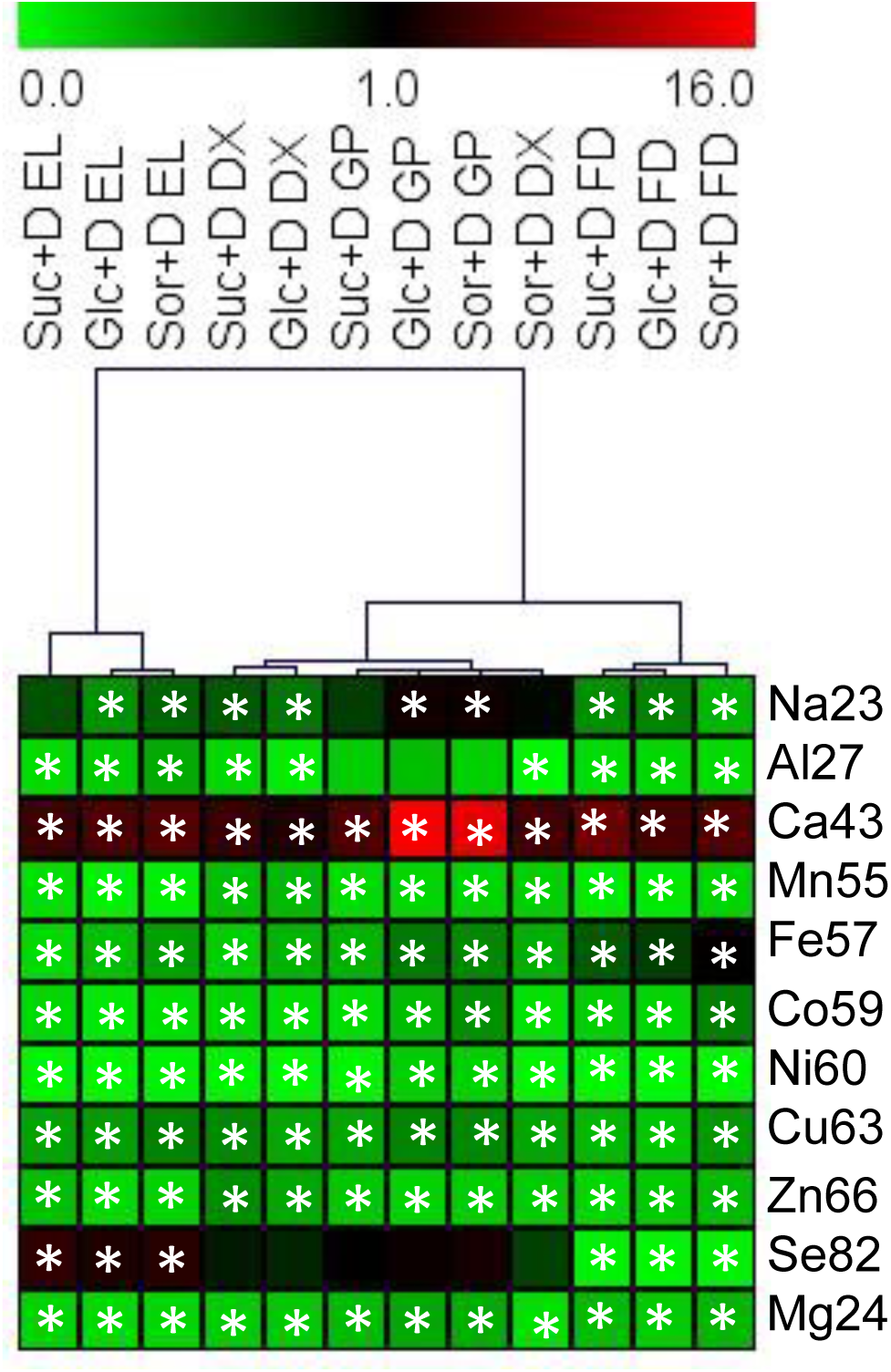
Heat map of showing the relative abundance of minerals in osmotic dehydrated peach slices with respect to fresh fruit. The colour scale is shown on the top of the figure. *denotes statistically significant differences of the mineral expressed in a DW basis in the OD+D with respect to the amount in fresh fruit (*p*<0.05). Gold Prince (GP), Elegant Lady (EL), Dixiland (DX) and Flordaking (FD). Hierarchical clustering analysis (HCA) was performed to group the samples using MeV4 software.

Hierarchical clustering (HCA) analysis was performed to group the samples. Interestingly, samples from the same cv group together, with those from EL forming a separate branch of the rest of the cvs (Fig. 5). Therefore, the changes in the mineral composition depend on the type of cv used rather than the osmolytes used for dehydration.

#### 3.2.6. Hierarchical clustering analysis reveals that OD+D are grouped depending on the cultivar

Hierarchical clustering analysis (HCA) of F, D and OD+D based on the complete data set obtained in this work was used to group similar samples into clusters (Fig. 6). As expected, F samples cluster together in a branch separated from dried peach (D and OD+D). Additionally, D samples cluster together and separately from OD+D. The most interesting feature is the clusterization of OD+D slices; samples were grouped based on their genotype and not by the HS used for pre-treatment. This categorization is in line with Two-way ANOVAs performed for each parameter (Table 3), which showed that in OD+D samples, MC, a*, L, AC, AsA, carotenoids, TPC, flavonoids, proteins, inhibition of AMY and protease, and inhibition of the growth of *B. cereus, S. aureus, E. faecalis* and *S. typhimurium* are not dependent on the HS used. In addition, in OD, neither ML, WL nor SG depended on the HS, in the conditions assayed in this work. Taken together, the choice of the cv is key in the properties of the resulting dehydrated peach, which influences in the colour, AC (DPPH), AsA, carotenoid, TPC, tannins and inhibition of the growth of all the testes microorganisms besides *E. faecalis* (Table 3).

**Fig. 6.**
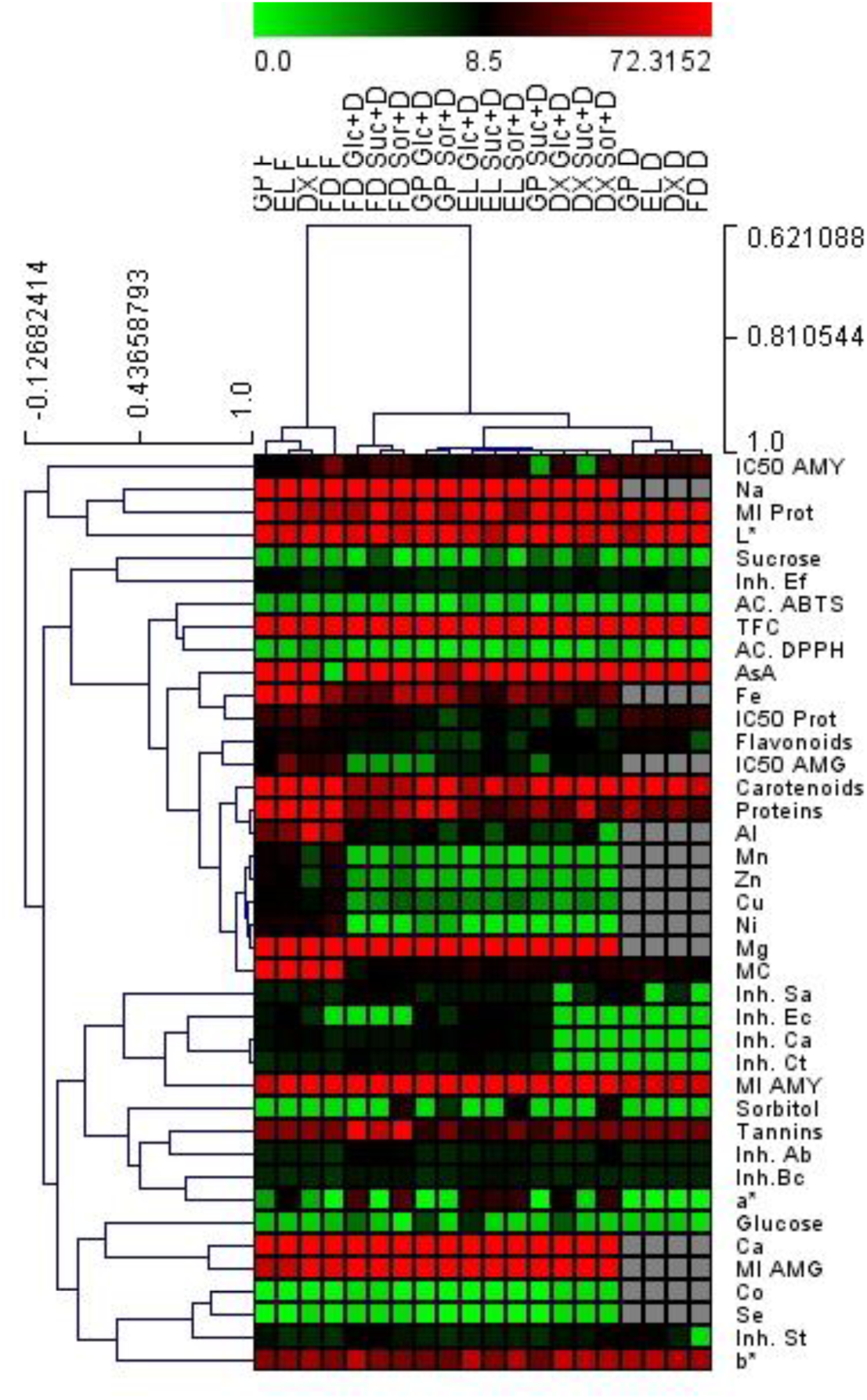
Hierarchical clustering analysis (HCA) of fresh (F), conventional dried (D) and OD treated followed hot air drying using sucrose (Suc+D), glucose (Glc+D) or sorbitol (Sor+D). Gold Prince (GP), Elegant Lady (EL), Dixiland (DX) and Flordaking (FD). The colour scale is shown on the top of the figure. HCA was performed to group the samples using MeV4 software. Ab, *Acinetobacter baumannii;* Bc, *Bacillus cereus;* Ca, *Candida albicans,* Ct, *C. tropicalis.* Ec, *Escherichia coli;* Ef, *Enterobacter faecalis*; Inh, microorganism growth inhibition; MI, maximum inhibition; Prot, Protease; Sa, *Staphylococcus aureus;* St, *Salmonella typhimurium*.

Due to high accessibility and low cost, Suc is the preferred osmolyte, followed by Glc. Besides it can be claimed that these pretreatments enlarge the sugars in the OD+D peach slices, values of Glc in Glc+D are lower than those of other fruits rich in this sugar and those of Suc in Suc+D are in the similar range or slightly higher than those of Suc in mango and grapes. Glc content of dehydrated slices ranged between 636.06±55.3 and 1425.13±63.3 µmol/GFW (Supplementary Tables S2-5) in Glc+D, which are equivalent to 11.5 g/100g and 25.7g/100g. For reference, Glc in dates ranged from 19.76 to 83.08 g/100g (AlShwyeh & Almahasheer, 2022) and in grapes from 19.36 to 25.46 g/100g (Zhong et al., 2023). Suc in Suc+D ranged between 318.36±23.3 and 1320.7±52.1 µmol/GFW, which are equivalent to 10.9 and 45.2 g/100g. For comparison, Suc in grapes fluctuated from 2.90g/100g to 6.40g/100g (Zhong et al., 2023) and in mangos it oscillated between 1.7 and 10.5 g/100 g (Bello-Pérez et al., 2007). Thus, although Glc or Suc are up-taken by peach during OD, peach slices are a wealthy food.

A growing number of consumers are aware of the high consumption of free sugars; therefore, sugar alcohols such as sorbitol emerge as suitable alternative to sugars, besides their higher price. In OD pre-treatments, the utilization of sorbitol rendered a good retention of carotenoids and phenolic compounds in kiwifruit (Kroehnke et al., 2021). Most recently, the use of maltitol with Suc improved the cohesiveness and springiness of the dehydrated peach slices (Wang et al., 2025). Sorbitol, which is a natural sugar alcohol of peach, is also used as sweetener, bulking agent, humectant, sequestrant, stabilizer and thickener. It has been considered as a safe food additive (Grembecka, 2015). Since it has lower calories than its correspondent monosaccharide, and considering that Sor+D have similar properties than Glc+D and Suc+D peach slices (Figs. 4-6, Tables 1-3), its use in the HS for OD treatments could be a recommended option for consumers interested in the in the control of calories intake.

## 4. Conclusion

Osmotic pretreatment of peach is a mild process through which water is removed from the fruit and solids are impregnated. It is used to extend the fruit short shelf life and to reduce the time of hot-air exposure to complete the dehydration process. Our results demonstrate that compared to exclusively hot-air-dried (D) peaches, it is better to maintain organoleptic and functional properties of the fruit by minimizing adverse changes such as carotenoid decrease and even improving some features such as tannin content and antimicrobial activities. Moreover, the use of Calcium in the HS enriches the amount of this mineral to the levels of other sources rich in this nutrient. Dehydrated-peach slices can be consumed as a snack or used in the food industry; therefore, the process of OD+D adds value to a natural product.

Besides the extensive characterization of dehydrated peach, which also includes the first study of the antimicrobial and the inhibition of enzyme activities in OD+D peach, the value of this work lies in the comparison of the three osmotic agents (Suc, Sor and Glc) and four peach cultivars. Under the incubation conditions tested (47°Brix HS at 40°C for 2h), the use of Suc, Glc or Sor rendered equivalent results in mass transfer, bioactive compound, AC and antimicrobial activity. On the contrary, differences among samples relay on the cv employed. In our case, EL and DX, both mid-season peach exhibited the more similar responses. In this respect, overproduction of the fruit is linked to the mid-season, where most of the cultivars are adapted. Therefore, the production of dehydrated fruit is a plausible use of the fruit at that time of the year. Additionally, the dissemination of this type of food will encourage the consumption of fruits over the whole year and to replace the consumption of ultra-processed snacks. OD+D improves the accessibility of a peach-derived item which can eaten as snack with a sugar content equivalent to other fruits when Glc or Suc are used for dehydration or even lower, when Sor is employed. Dehydrated peach slices will thus attempt to reduce the risk of development of diverse diseases associated with inflammatory process, hyperglycemia and oxidative stress.

Research on sensorial characteristics and storage stability of osmotically dehydrated peach slices is needed and is currently being conducted. Moreover, the management of the HS, such as the re-use of the solution, is also being studied.

## CRediT authorship contribution statement

**Lara Salvañal:** Methodology, Investigation, Formal analysis, Visualization. **Julieta Gabilondo:** Methodology, Investigation, Writing – review & editing, Conceptualization. **Graciela Corbino:** Methodology, Investigation, Conceptualization. **Claudio O. Budde:** Methodology, Investigation, Conceptualization. **María Valeria Lara:** Writing – original draft, review & editing, Supervision, Resources, Project administration, Funding acquisition, Data curation, Visualization, Conceptualization.

## Data availability

Data will be made available on request.

## Appendix A. Abbreviation used

The Abbreviation used in the study are ABTS*^+^, 2,2-azinobis (3-ethylbenzothiazoline-6-sulfonic acid (ABTS*^+^);

AC: antioxidant capacity
AMG: α-amyloglucosidase
AMY: α-amylase
AsA: ascorbic acid
CA: chlorogenic acid equivalents
Cv: cultivar
D: hot air drying
DMSO: dimethyl sulfoxide
DW: dry weight
DPPH: 1,1-diphenyl-2-picrylhydrazyl (DPPH)
DX: Dixiland
EL: Elegant Lady
F: Fresh fruit
FD: Flordaking
FW: fresh weight
GDW: gram of dry weight
GFW: gram of fresh weight
Glc: glucose
GP: Goldprince
HS: hyperosmotic solution
ICP-MS: inductively coupled plasma-mass spectrometry
MC: moisture content
ML: mass loss
OD: osmotic dehydration
SG: solute gain
Sor: sorbitol
Suc: sucrose
RE: rutin equivalents
TA: titratable acidity
TCA: trichloroacetic acid
TE: Trolox equivalents
TPC: total phenolics
Trolox: 6-hydroxy-2, 5, 7, 8-tetramethyl chromene-2-carboxylic acid
TSS: total soluble solids
WL: water loss.

## Supplementary data

**Supplementary Table S1.** Mineral quantification in fresh (F) and OD-D peach slices. ICP-MS was conducted in digested and filtered samples. Five biological replicates were used. Slices were incubated in HSs of Suc, Glc or Sor and the hot air dried (Suc+D, Glc+D or Sor+D, respectively). Total mineral concentrations were expressed as µg per GDW.

| Cultivar | Element | Sample |  |  |  |  |  |  |  |
| --- | --- | --- | --- | --- | --- | --- | --- | --- | --- |
|  |  | Fresh (µg/GDW) |  | Suc+D (µg/GDW) |  | Glc+D (µg/GDW) |  | Sor+D (µg/GDW) |  |
|  |  | Mean | SD | Mean | SD | Mean | SD | Mean | SD |
| Gold Prince | Na 23 | 420.464241 | 20.8642657 | 320.259276 | 132.138184 | 865.569718 | 126.280033 | 819.231424 | 147.397222 |
| Gold Prince | Al 27 | 33.5799999 | 4.17704998 | 6.56154161 | 2.05724237 | 8.63457278 | 0.45618813 | 6.20994594 | 1.78636894 |
| Gold Prince | Ca 43 | 403.95416 | 98.9314343 | 2219.9378 | 318.681961 | 6306.79752 | 1132.22566 | 5834.19106 | 701.805686 |
| Gold Prince | Mn 55 | 11.621469 | 0.61342657 | 1.64396967 | 0.16077631 | 1.84993532 | 0.30301323 | 1.92904439 | 0.63300772 |
| Gold Prince | Fe 57 | 114.068015 | 10.2212783 | 34.0460605 | 3.22694951 | 59.9192304 | 15.8083984 | 52.664145 | 5.27829538 |
| Gold Prince | Co 59 | 0.18005183 | 0.02972546 | 0.02296367 | 0.00450434 | 0.04639821 | 0.00763399 | 0.07792261 | 0.06592535 |
| Gold Prince | Ni 60 | 13.174559 | 1.70414638 | 0.55181098 | 0.16105009 | 2.49467129 | 0.29163162 | 2.46266724 | 0.13728949 |
| Gold Prince | Cu 63 | 9.6052737 | 0.30213354 | 3.3267929 | 0.29125248 | 4.56871116 | 0.98086948 | 4.49156006 | 1.19900078 |
| Gold Prince | Zn 66 | 11.7903188 | 4.37962549 | 2.55715085 | 0.24275116 | 1.95338812 | 0.47727158 | 2.34619819 | 0.46960438 |
| Gold Prince | Se 82 | 0.02370255 | 0.01327579 | 0.02664065 | 0.0083501 | 0.04117568 | 0.00860968 | 0.05331055 | 0.04216238 |
| Gold Prince | Mg 24 | 762.088059 | 127.023941 | 206.503294 | 47.0083544 | 255.010325 | 39.7450504 | 247.726909 | 67.1250682 |
| Elegant Lady | Na 23 | 475.0649 | 57.8801 | 366.256292 | 17.7097633 | 195.910508 | 29.9030104 | 275.86349 | 51.050329 |
| Elegant Lady | Al 27 | 41.8601519 | 1.36397004 | 5.73264471 | 0.31463453 | 8.18513115 | 1.22022868 | 13.5985139 | 3.30475441 |
| Elegant Lady | Ca 43 | 723.483198 | 72.6639299 | 3384.08871 | 794.841155 | 3926.1846 | 517.630629 | 4288.44988 | 328.19214 |
| Elegant Lady | Mn 55 | 14.0909706 | 0.60515023 | 1.75204019 | 0.0497895 | 0.81850739 | 0.13144388 | 0.84434838 | 0.11612074 |
| Elegant Lady | Fe 57 | 124.244843 | 4.46125815 | 18.4273611 | 4.23683805 | 29.8669783 | 7.02182115 | 46.2404212 | 15.8996085 |
| Elegant Lady | Co 59 | 0.21750205 | 0.010055 | 0.03196171 | 0.00084793 | 0.01820046 | 0.00393154 | 0.02611473 | 0.00317835 |
| Elegant Lady | Ni 60 | 13.989051 | 1.13422055 | 0.60574103 | 0.05495608 | 0.51871313 | 0.11487165 | 0.69273503 | 0.06012564 |
| Elegant Lady | Cu 63 | 10.6113791 | 0.50944081 | 3.80454019 | 0.5884372 | 4.02879589 | 1.09046992 | 5.084761 | 0.51804332 |
| Elegant Lady | Zn 66 | 10.2507491 | 0.43639308 | 2.55715085 | 0.24275116 | 1.4803288 | 0.3904694 | 1.73332786 | 0.60164409 |
| Elegant Lady | Se 82 | 0.00664904 | 0.0000001 | 0.02545673 | 0.00335103 | 0.01862478 | 0.00422986 | 0.02282534 | 0.00266837 |
| Elegant Lady | Mg 24 | 837.134906 | 31.1842535 | 149.174913 | 24.6315742 | 124.445118 | 26.6007481 | 133.304799 | 25.6247054 |
| Dixiland | Na 23 | 436.664463 | 45.7942174 | 283.266961 | 56.7896968 | 231.965117 | 51.1923483 | 423.959518 | 18.4961512 |
| Dixiland | Al 27 | 88.8954671 | 18.0607474 | 12.9057204 | 12.1528667 | 6.3459529 | 3.73093948 | 1.89326579 | 0.201739 |
| Dixiland | Ca 43 | 628.957985 | 94.0956543 | 2666.95384 | 1105.78943 | 1789.08113 | 320.357583 | 3038.90756 | 213.297742 |
| Dixiland | Mn 55 | 6.19264283 | 0.90262391 | 1.53642051 | 0.2904405 | 1.68558116 | 0.12974325 | 1.1504438 | 0.14921489 |
| Dixiland | Fe 57 | 108.377697 | 13.0472932 | 21.2416938 | 3.07775249 | 31.935711 | 5.49015174 | 33.1677128 | 9.6362671 |
| Dixiland | Co 59 | 0.17026971 | 0.01326802 | 0.02771703 | 0.0050658 | 0.02213429 | 0.00901486 | 0.01807917 | 0.00419271 |
| Dixiland | Ni 60 | 12.5630206 | 1.24490995 | 0.61330572 | 0.05438585 | 0.45767041 | 0.08782979 | 0.54195019 | 0.06746645 |
| Dixiland | Cu 63 | 8.10779592 | 0.90571241 | 3.76514438 | 0.27428479 | 2.73459997 | 0.16454204 | 2.82353599 | 0.28827323 |
| Dixiland | Zn 66 | 5.69223845 | 0.14053933 | 2.58511217 | 0.22445696 | 1.97001686 | 0.44488564 | 1.15637051 | 0.30366511 |
| Dixiland | Se 82 | 0.02732116 | 0.01381057 | 0.0245487 | 0.00206398 | 0.02342934 | 0.00859492 | 0.02043998 | 0.00183172 |
| Dixiland | Mg 24 | 815.798849 | 116.350221 | 167.430381 | 62.2724555 | 175.798238 | 34.6631468 | 104.481322 | 22.3137031 |
| Flordaking | Na 23 | 424.247768 | 54.4747513 | 187.707328 | 12.5185525 | 205.101547 | 33.5909689 | 121.060519 | 23.034168 |
| Flordaking | Al 27 | 56.1961012 | 9.50588208 | 7.75312856 | 0.52147145 | 8.35834338 | 1.56335202 | 7.78505487 | 2.87868894 |
| Flordaking | Ca 43 | 455.985728 | 94.804628 | 3313.02394 | 76.7762981 | 2350.90097 | 427.787624 | 2654.48885 | 152.340343 |
| Flordaking | Mn 55 | 20.0111515 | 0.76915799 | 1.68205839 | 0.12705112 | 1.71811539 | 0.55901064 | 3.31242452 | 0.64817201 |
| Flordaking | Fe 57 | 45.5048623 | 1.94307746 | 29.4040162 | 0.39797625 | 29.3320741 | 7.70954224 | 57.5779179 | 4.87694855 |
| Flordaking | Co 59 | 0.1199322 | 0.03703125 | 0.02191398 | 0.00307181 | 0.01899982 | 0.00437278 | 0.05915616 | 0.01585819 |
| Flordaking | Ni 60 | 21.7520096 | 1.25662364 | 0.70181242 | 0.06661969 | 0.47871803 | 0.10768275 | 0.80355659 | 0.14933818 |
| Flordaking | Cu 63 | 13.6451152 | 2.27889547 | 3.79526232 | 0.38274787 | 2.93336706 | 0.63534677 | 5.396891 | 1.24301087 |
| Flordaking | Zn 66 | 15.3052088 | 5.21761891 | 2.45356121 | 0.14344991 | 3.16033872 | 0.9944136 | 4.05480328 | 0.85771619 |
| Flordaking | Se 82 | 0.31668584 | 0.02524757 | 0.0149554 | 0.00172871 | 0.01106516 | 0.00266676 | 0.01578627 | 0.00172684 |
| Flordaking | Mg 24 | 1046.39514 | 11.1604205 | 271.794687 | 22.0762877 | 205.214944 | 54.0703842 | 298.286085 | 7.11106388 |

**Supplementary Table S2.** **ΔE*76** parameter of peach slices from hot air dried (D) or OD treated slices (Suc, Glc, Sor) followed by hot air drying (Suc+D, Glc+D, Sor+D) with respect to fresh fruit (F). Gold Prince (GP), Elegant Lady (EL), Dixiland (DX) and Flordaking (FD). Scale for classification is shown at the bottom of the table.

| | $\Delta E^*76$ | | | |
| --- | --- | --- | --- | --- |
|  | GP | EL | DX | FD |
| Suc | 11.83 | 8.29 | 8.52 | 12.29 |
| Suc+D | 10.31 | 15.68 | 9.50 | 4.32 |
| Glc | 14.00 | 8.14 | 19.43 | 15.71 |
| Glc+D | 8.69 | 15.72 | 13.55 | 37.21 |
| Sor | 13.13 | 18.18 | 28.85 | 32.28 |
| Sor+D | 10.19 | 16.42 | 19.78 | 34.53 |
| D | 17.57 | 17.52 | 6.78 | 11.06 |

**Not visible:** $\Delta E^*76 < 0.2$
**Very low:** $0.2 \leq \Delta E^*76 \leq 0.5$
**Low:** $0.5 < \Delta E^*76 \leq 1.5$
**Noticeable:** $1.5 < \Delta E^*76 \leq 3.0$
**Very noticeable:** $3.0 < \Delta E^*76 \leq 6.0$
**High:** $6.0 < \Delta E^*76 \leq 12.0$
**Very High:** $\Delta E^*76 > 12.0$

**Supplementary Fig. S1.**
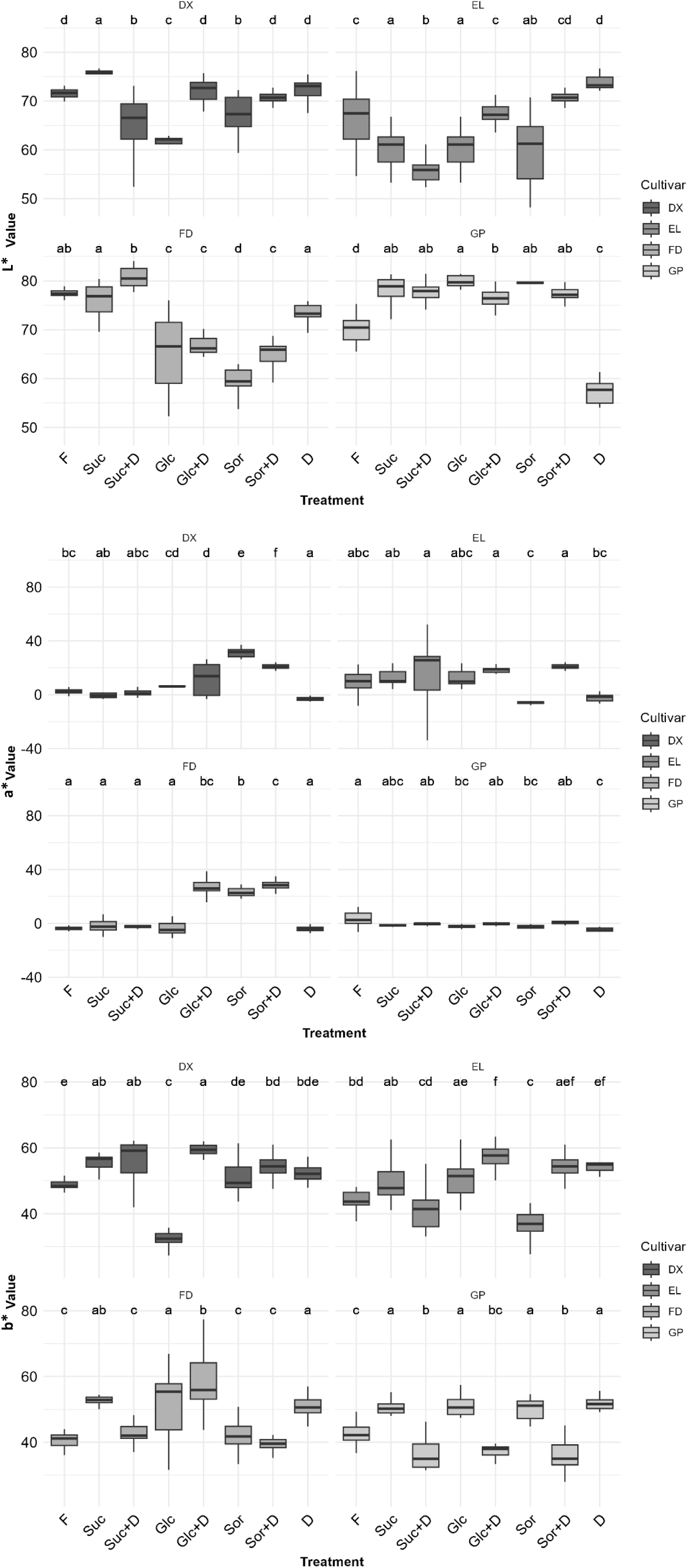
Colour parameters of peach slices from fresh (F), hot air dried (D) or OD treated slices (Suc, Glc, Sor) followed by hot air drying (Suc+D, Glc+D, Sor+D). Gold Prince (GP), Elegant Lady (EL), Dixiland (DX) and Flordaking (FD). Bars represent the mean of five replicates ± SD. Letters on top of each figure show the statistical analysis. Values with at least one same letter are not statistically significant according to ANOVA followed by Tukey test, *p ≤* 0.05.

**Supplementary Fig. S2.**
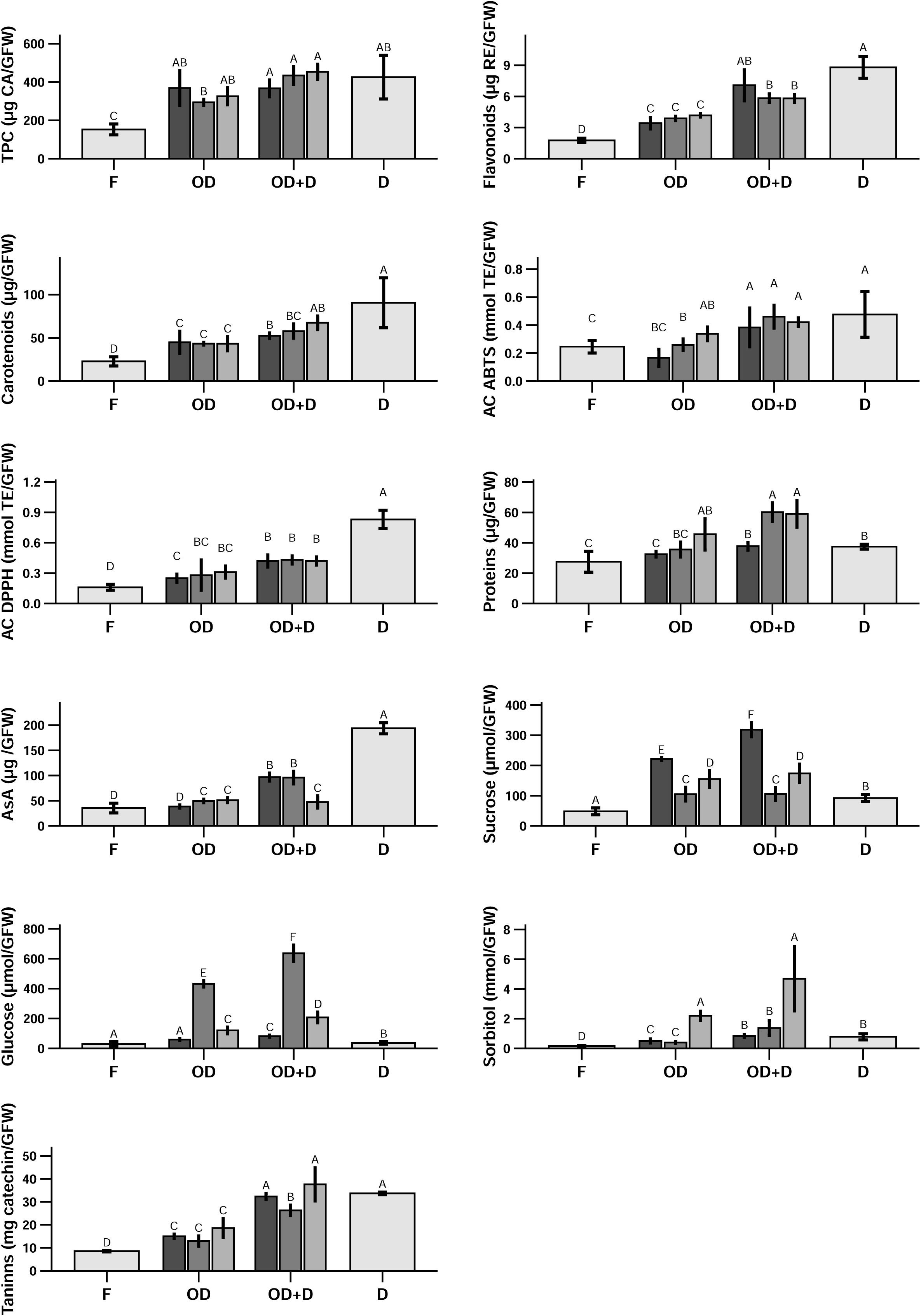
Bioactive compound contents, antioxidant capacity (AC) and nutritional properties of Gold Prince (GP) fresh peach fruit (F), dried slices (D), or slices incubated with HSs of Suc, Glc or Sor (OD) followed by hot air drying (OD+D). Bars with at least one same letter are not statistically significant different (ANOVA, Tukey test, *p ≤* 0.05). Results are expressed in a GFW basis.

**Supplementary Fig. S3.**
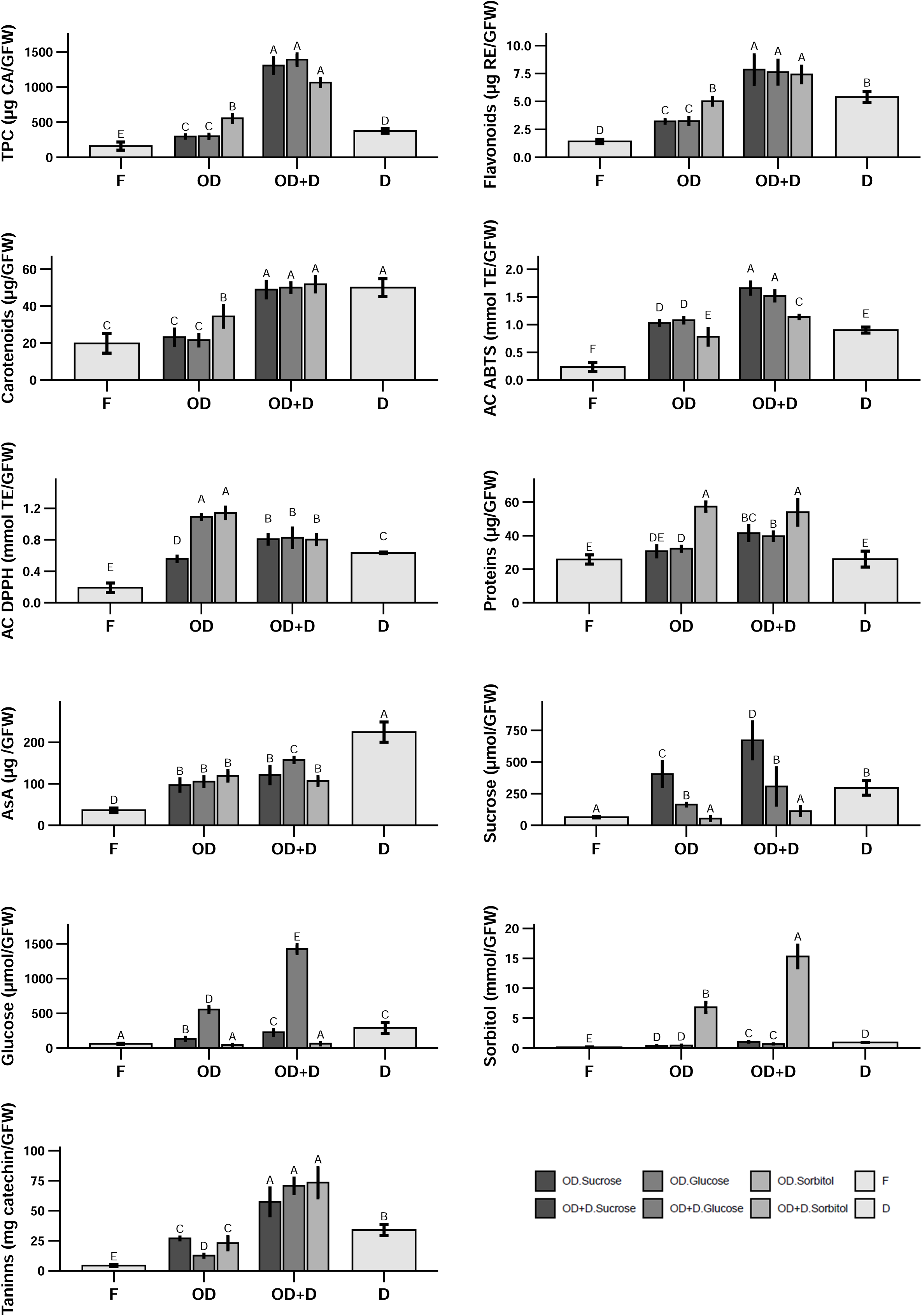
Bioactive compound contents, antioxidant capacity (AC) and nutritional properties of Elegant Lady (EL) fresh peach fruit (F), dried slices (D), or slices incubated with HSs of Suc, Glc or Sor (OD) followed by hot air drying (OD+D). Bars with at least one same letter are not statistically significant different (ANOVA, Tukey test, *p ≤* 0.05). Results are expressed in a GFW basis.

**Supplementary Fig. S4.**
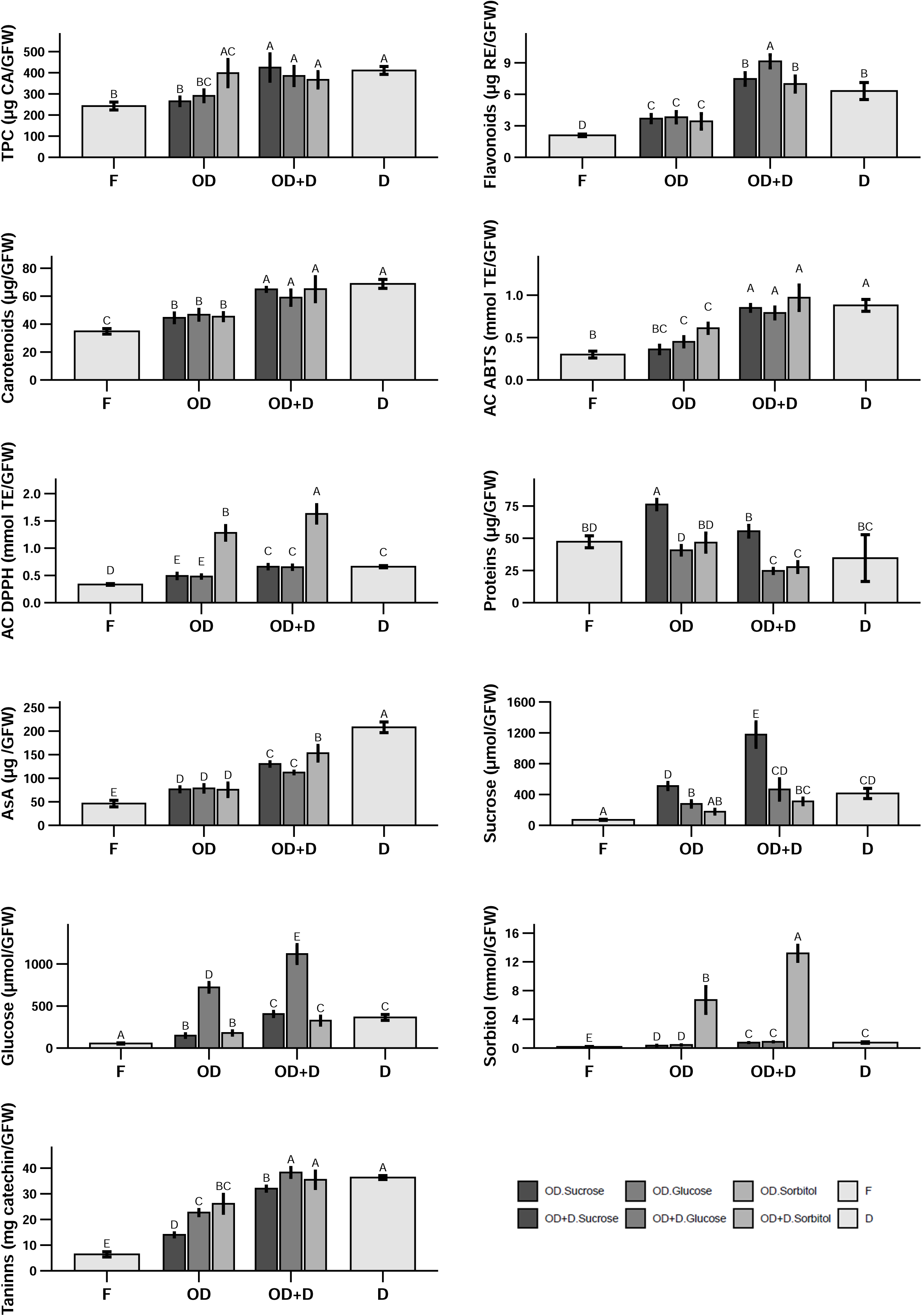
Bioactive compound contents, antioxidant capacity (AC) and nutritional properties of Dixiland (DX) fresh peach fruit (F), dried slices (D), or slices incubated with HSs of Suc, Glc or Sor (OD) followed by hot air drying (OD+D). Bars with at least one same letter are not statistically significant different (ANOVA, Tukey test, *p ≤* 0.05). Results are expressed in a GFW basis.

**Supplementary Fig. S5.**
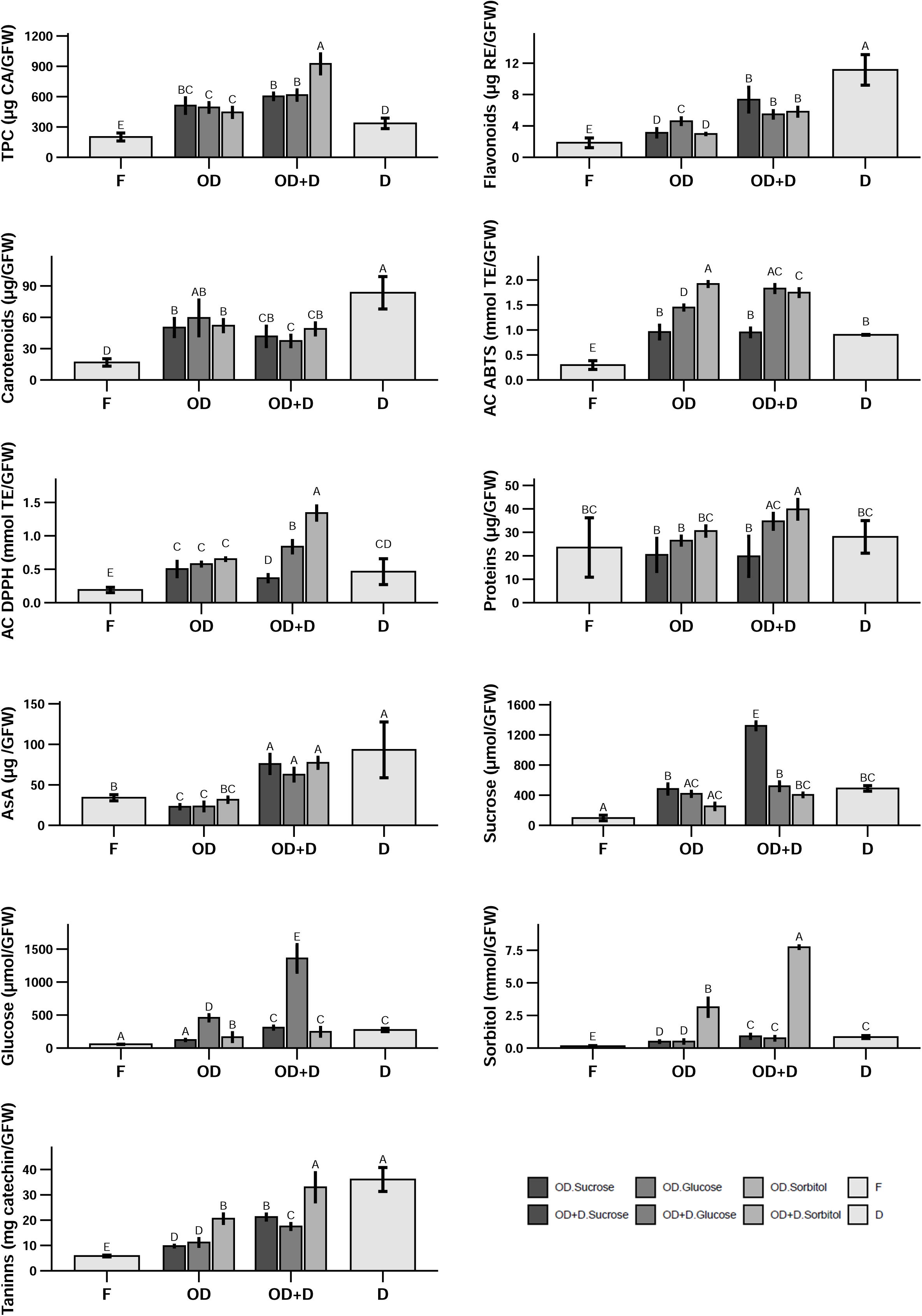
Bioactive compound contents, antioxidant capacity (AC) and nutritional properties of Flordaking (FD) fresh peach fruit (F), dried slices (D), or slices incubated with HSs of Suc, Glc or Sor (OD) followed by hot air drying (OD+D). Bars with at least one same letter are not statistically significant different (ANOVA, Tukey test, *p ≤* 0.05). Results are expressed in a GFW basis.

